# The temporal dynamics of animacy-related neural representations are robust to controversial training

**DOI:** 10.64898/2026.11.12.766140

**Authors:** Leylan Öykü Etkin, Enes Bahadır Kömürcü, Chiu-Yueh Chen, Hans Op de Beeck, Andrea Ivan Costantino

## Abstract

Animal bias refers to the human tendency to prioritise the neural processing of animate over inanimate stimuli, a phenomenon observed across multiple cognitive domains, such as attention, perception, memory, and language. Lookalikes (also referred to as zoomorphic objects), inanimate objects that visually resemble animals, tend to be interpreted as animate regardless of their inanimate identity. We examined whether this animal bias can be modified through learning to understand how experience and plasticity shape visual object representations. We tested whether feedback-based categorisation training in which lookalikes had to be categorised as inanimate could reduce the animal bias. Thirty healthy participants completed three weekly sessions of animacy categorisation training, followed by an EEG recording session during which they performed an animacy task. Repeated-measures ANOVA on behavioural data revealed that categorisation efficiency, measured as the inverse efficiency score, improved both within and across sessions, but there was no evidence that this improvement was specific to lookalikes. Time-resolved multivariate pattern analysis and representational similarity analysis on EEG data obtained after this training showed a descriptively smaller animal bias relative to the no-training dataset from Chen et al. (2023), but this reduction was not statistically significant in any analysis. Moreover, a significant animal bias was still present in participants who underwent training. Thus, neural reorganisation related to the animal bias is limited even after several training sessions to overcome this bias.

## 1 Introduction

Visual object recognition allows humans to identify objects rapidly and accurately, even under substantial changes in their appearance (DiCarlo et al., 2012; Maniquet et al., 2025). A central neural pathway supporting this ability is the ventral visual stream (VVS), in which objects are represented through distributed patterns of neural activity (Bracci & Op de Beeck, 2023; Kriegeskorte et al., 2008). One of the most prominent organising dimensions of these representations is animacy, the distinction between animate and inanimate objects (Bracci & Op de Beeck, 2016; Grill-Spector & Weiner, 2014; Konkle & Caramazza, 2013; Kriegeskorte et al., 2008; Naselaris et al., 2012; Ritchie et al., 2021). Category-selective areas, including regions responsive to faces, bodies, places and written words, have long provided evidence for the organisation of the VVS (Cohen et al., 2000, 2002; Downing et al., 2001; Epstein & Kanwisher, 1998; Kanwisher et al., 1997), although the extent to which such selectivity reflects object category rather than shared visual features remains debated (Bracci et al., 2017; Pillet et al., 2024). The distinction between animate and inanimate stimuli is also relevant outside vision, including attention and memory, and sensitivity to animacy emerges early in development (Hofrichter et al., 2021; Nairne et al., 2013; Rakison & Poulin-Dubois, 2001; Serra & DeYoung, 2023; Serra et al., 2024). Even simple moving shapes can elicit the perception of animate agents and the attribution of mental states (Gergely & Csibra, 2003; Gergely et al., 1995; Heider & Simmel, 1944), a capacity known as theory of mind (Premack & Woodruff, 1978), emphasising that animacy judgements are not determined exclusively by an object’s physical identity. Visual appearance and an object’s actual animacy do not always agree. Lookalikes—inanimate objects that visually resemble animals—provide a particularly informative case because they contain animal-like features despite their inanimate identity.

Features such as global shape, curvature, and eye- or mouth-like configurations can provide cues to animacy (Bracci et al., 2019; Kato & Mugitani, 2015; Keefe et al., 2018; Long et al., 2017, 2018; Wardle et al., 2020). For ordinary objects, such cues often support efficient categorisation. In lookalikes, however, the same features may compete with knowledge of the object’s true identity, making them an informative test of how appearance and categorical knowledge shape neural representations. Using fMRI and multivariate pattern analysis (MVPA), Bracci et al. (2019) showed that although observers categorise lookalikes as inanimate, their neural representations in the VVS are more similar to those of real animals than to those of other inanimate objects. Representational similarity analysis (RSA) supported the same pattern, showing an appearance-driven organisation of lookalikes in neural representational space. We refer to this greater neural similarity of lookalikes to animals as the *animal bias*. Related EEG and MEG research has shown that shape and mid-level visual features contribute substantially to neural distinctions between animate and inanimate objects, although they may not fully account for them (Grootswagers et al., 2019; Kaiser et al., 2016; Proklova et al., 2019). These findings make it important to distinguish the representation of animal-like appearance from knowledge of an object’s inanimate identity (Jozwik et al., 2022).

The animal bias is also expressed in the temporal dynamics of visual processing. Using time-resolved EEG decoding, Chen et al. (2023) found that lookalikes initially evoke neural response patterns more similar to those of real animals, whereas their inanimate identity becomes more evident at later stages. This progression is consistent with evidence that early object responses are strongly influenced by visual features, while later responses increasingly reflect category-related information (T. Carlson et al., 2013; Cichy et al., 2014; Contini et al., 2017), a sequence also observed in attentional selection during visual search (Yeh & Peelen, 2022). In Chen et al. (2023), fMRI–EEG fusion analyses further associated early animal-bias representations with posterior regions of the VVS and later processing with more anterior regions. This temporal profile suggests that the animal bias is expressed during relatively early visual processing, rather than appearing only during explicit object categorisation. Whether these early neural representations can be modified through experience remains unclear.

Comparisons with deep neural networks (DNNs), which predict ventral stream responses well (Cichy et al., 2016; Khaligh-Razavi & Kriegeskorte, 2014; Yamins et al., 2014), provide a possible link between the animal bias and visual experience. Standard object-classification networks do not typically reproduce the human tendency to represent lookalikes together with animals (Bracci et al., 2019). However, this bias can emerge when DNNs are explicitly trained to group lookalikes with animals, producing representational patterns more similar to those observed in humans (Duyck, Costantino et al., 2024). This finding suggests that learned visual statistics may contribute to the human animal bias. Extensive exposure to animal-like shapes could, in principle, help establish neural representations that are difficult to reshape in adulthood, but the DNN findings do not determine whether this is the case in humans. An important question therefore follows: can targeted experience modify an already established bias in the adult visual system, or is it relatively resistant to short-term training?

The adult visual cortex retains a capacity for experience-dependent plasticity (Bang et al., 2023; Cheng et al., 2025; Watanabe & Sasaki, 2015), and targeted training can alter visual object representations (Baeck et al., 2016; Mayhew et al., 2010). Category learning has been proposed to reshape representational geometry by increasing the separation of stimuli along task-relevant dimensions (*dimensional modulation*; Folstein et al., 2015). In this account, training changes the weight assigned to diagnostic features, making stimuli from competing categories more distinguishable. Learning semantic associations can similarly alter representations of novel objects in the VVS (Clarke et al., 2016). Moreover, changes in perceptual representations may be especially pronounced for ambiguous stimuli near category boundaries, where identifying relevant features is particularly demanding (O’Bryan et al., 2024). Lookalikes are analogous to such ambiguous exemplars because their animal-like appearance conflicts with their inanimate identity. However, previous demonstrations of plasticity have often involved simple or novel stimuli; it remains unknown whether similar training can modify a bias associated with familiar animal-like features and extensive visual experience.

Here, we tested whether multi-session, feedback-based categorisation training could reduce the neural animal bias. Participants were trained to categorise lookalikes according to their inanimate identity rather than their animal-like appearance, a task designed to encourage attention to features diagnostic of the correct category (Van Gulick & Gauthier, 2014). Training included many exemplars to support learning beyond individual images, given that visual learning can otherwise remain specific to the trained stimuli or viewpoints (Baeck et al., 2012, 2016). Importantly, the stimuli used in the subsequent EEG test were not presented during training, allowing us to investigate whether any effects generalised to untrained exemplars. The EEG task required participants to categorise lookalikes as inanimate, so the test measured the neural organisation of objects under an explicit identity-based categorisation task. To assess neural representations over time, we used the animacy task and time-resolved EEG approach of Chen et al. (2023) and compared the trained participants with their previously published untrained sample.

We hypothesised that feedback-based categorisation training would improve performance for lookalikes across sessions (H1), reflecting greater sensitivity to their inanimate identity. The training was “controversial” in the sense that it required responses based on object identity despite conflicting animal-like appearance, by analogy with “controversial stimuli” used to distinguish between computational models (Golan et al., 2020). We expected less improvement for animals and other inanimate objects, for which appearance and identity are less ambiguous. Behavioural performance was measured using the inverse efficiency score (IES), combining response time and accuracy (Townsend & Ashby, 1978).

Based on Chen et al. (2023), we expected animacy information to become decodable early in the EEG response (approximately 80 ms after stimulus onset). For lookalikes, we considered two possible outcomes following training (H2), assessed by comparing their neural decoding patterns with those from the untrained sample of Chen et al. (2023). If the established bias can be modified, lookalikes should become neurally more distinguishable from animals and more similar to inanimate objects, reflected in a reduction in the difference between lookalike–inanimate and lookalike–animate decoding accuracies, particularly around 100–400 ms after stimulus onset. Alternatively, if the bias is relatively resistant to short-term training, this difference should remain comparable to that in the untrained sample. Such persistence would be consistent with shared representational mechanisms and extensive visual experience. These predictions concern the neural separability of lookalikes from the two other categories, rather than changes in behavioural categorisation alone (Figure 1a).

**Figure 1.**
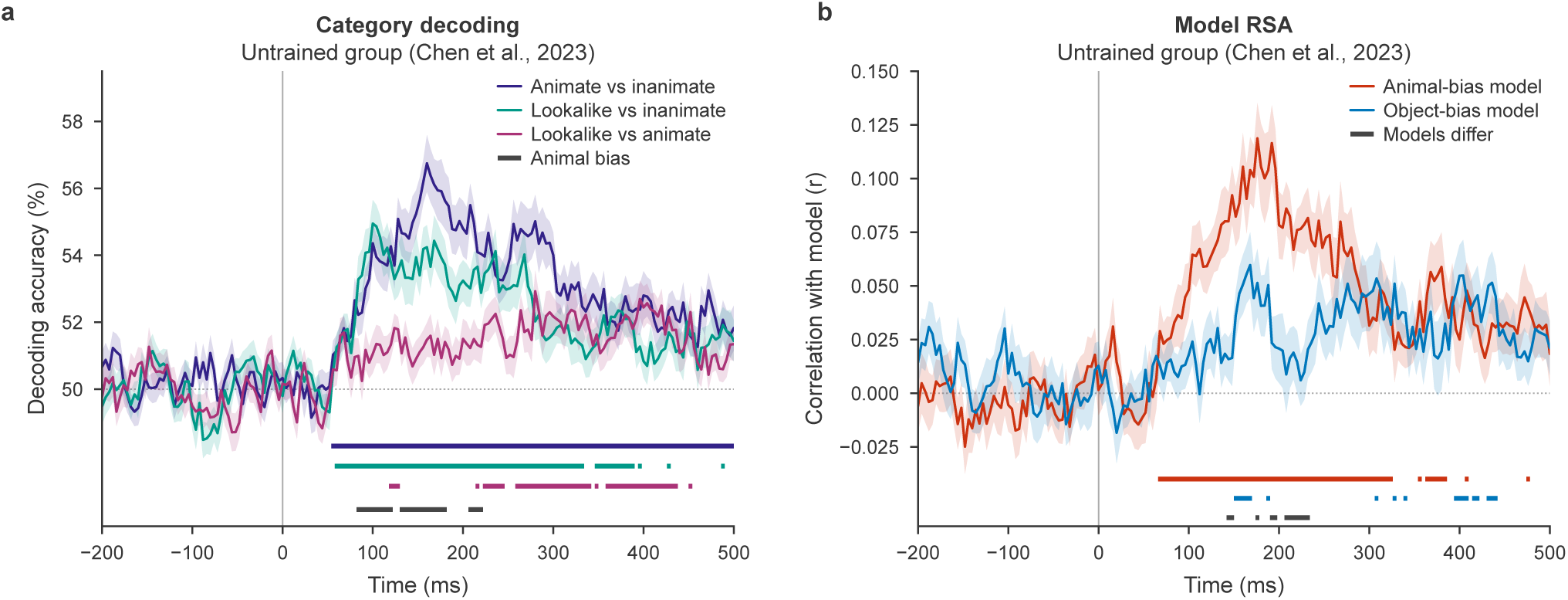
Predictions based on the untrained group of Chen et al. (2023). (a) Time-resolved pairwise decoding accuracy for three category comparisons. Coloured bars indicate time points of significant above-chance decoding, and the dark bar indicates a significant animal bias: higher lookalike–inanimate than lookalike–animate decoding accuracy. H2 predicts a reduction in this bias if training modifies lookalike representations. (b) Time-resolved correlations between EEG representational dissimilarity matrices (RDMs) and animal bias or object bias models. Coloured bars indicate significant model correlations, and the dark bar indicates a significant difference between models. H3 predicts reduced correspondence with the animal bias model relative to the object bias model following training. Lines show group means and shaded areas the standard error of the mean; the dotted line indicates chance (50%) or zero, and the vertical line indicates stimulus onset. Significance bars: *p* < .05, TFCE-corrected.

Finally, we hypothesised that training-related changes would be especially evident for lookalikes, whose appearance and identity conflict (O’Bryan et al., 2024). Using RSA, we tested whether training shifted the neural geometry of lookalikes toward an *object bias model*, which groups them with inanimate objects, and away from an *animal bias model*, which groups them with animals (H3; Figure 1b).

## 2 Methods

### 2.1 Participants

Thirty healthy individuals (24 females, 6 males) aged between 18 and 45 (*M* = 24.33, *SD* = 5.86) completed all three sessions of the study and were included in the final sample. All participants were right-handed and had normal or corrected-to-normal eyesight. The study was approved by the Social and Societal Ethics Committee of KU Leuven (G-2024-8866-R2(MIN)). Participants were recruited online via the Experimental Management System (Sona Systems) of KU Leuven. Before starting the experiment, all participants signed the information sheet and informed consent. Recruitment continued until 30 participants had completed all three sessions. In total, 36 individuals were initially recruited, of whom 30 completed all three sessions and the remaining 6 withdrew before completing all parts of the study. Participants who withdrew before completing all sessions were compensated on a pro-rated, per-session basis for the sessions they attended. Those who did not participate in any session received no compensation. Upon completion of all parts of the study, participants received either monetary compensation (60 euros) or course credits (4 credits). These characteristics of the tested sample are very similar to the sample in Chen et al. (2023).

### 2.2 Procedure

Participants completed three sessions in total over three weeks. The first two sessions were training sessions, during which participants practised discriminating animals (animate) from lookalikes and inanimate objects (both inanimate). The third session was the test session and included two parts: participants first completed the training task one last time, followed by an EEG session as the testing phase adapted from an earlier study (Chen et al., 2023).

### 2.3 Training phase

#### 2.3.1 Stimulus Set

The task for the training phase was programmed in PsychoPy, version 2024.2.4 (Peirce et al., 2019). The initial stimulus set included 520 different greyscale images representing three categories—animals (210), lookalikes (155), and inanimate objects (155). These images were the stimulus set used to fine-tune the object bias and animal bias deep neural networks in Duyck et al. (2024). The stimuli were presented to participants on a Dell screen (21.5 inches, 60 Hz, 1920 *×* 1080 pixel resolution). Because original image sizes varied, the size of each stimulus was standardised to 300 *×* 400 pixels in PsychoPy. A fixed set of 100 images (50 animals, 25 lookalikes, and 25 inanimate objects) was created from the initial stimulus set and presented twice during the task, once before the main training blocks (extra before) and once after (extra after), resulting in a total of 620 trials per participant (260 animal, 180 lookalike, and 180 inanimate object trials). For the practice block, a separate set of 10 cartoon-like images of animals, lookalikes, and inanimate objects was used.

#### 2.3.2 Task Description

Participants were asked to categorise the images based on their animacy as quickly and as accurately as possible. To complete this task, participants pressed either the left or the right arrow keys to indicate whether the stimulus presented was animate (animal) or inanimate (lookalikes and inanimate objects). To prevent motor response confounds, the key-category mappings were alternated between each block.

A training session consisted of a practice block and 6 training blocks, lasting for 25–30 minutes (Figure 2a). All the stimuli in the practice block had a display time of 1,000 ms. The extra before and extra after training blocks each contained 100 trials selected from the pool of 520 images, where the response time and accuracy did not affect the display time of the stimulus. For these blocks, the display time was always 600 ms. These blocks with a fixed stimulus duration were used to compare performance across blocks and sessions. For the remaining four training blocks in between, which contained 420 trials (105 trials per block), the stimulus display time started from 1,000 ms and adapted dynamically based on the accuracy and speed of participants’ responses. If the participant maintained 80% accuracy and responded faster than 70% of the stimulus display time during the last five trials, the display time of the following stimulus decreased by 5%. The stimulus display time never goes below 300 ms, no matter how accurate or fast the participants responded. In the same way, if the response accuracy dropped below 50% and the response time was slow relative to the stimulus duration, the subsequent stimulus duration increased by 5%. We included these blocks with an adaptive stimulus duration to increase the odds that training would also affect the initial processing phases.

**Figure 2.**
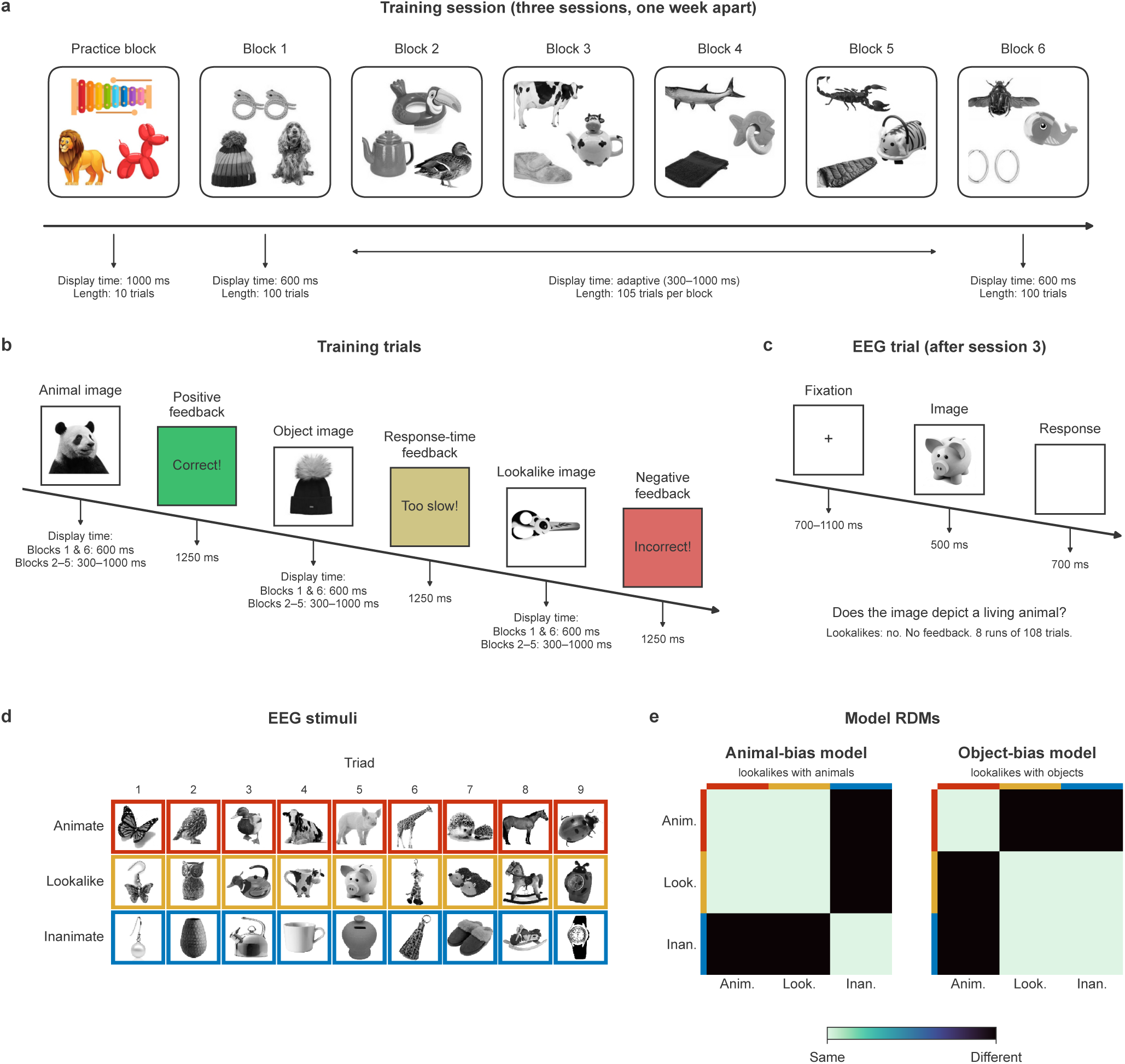
Experimental design. (a) One training session: a practice block (10 trials, 1,000 ms display time) followed by Blocks 1–6. Blocks 1 and 6 (extra before and extra after; 100 trials each) had a fixed display time of 600 ms, and Blocks 2–5 (105 trials each) a display time that adapted to performance between 300 and 1,000 ms. Participants completed three sessions, one week apart. Tiles show example training images. (b) Training trials: each image was followed by feedback for 1250 ms, positive (green), response-time (yellow) or negative (red). (c) EEG trial: a fixation cross (700–1100 ms), the image (500 ms) and a 700 ms response window, without feedback. Participants reported whether the image depicted a living animal; lookalikes required the inanimate response. (d) The 27 EEG stimuli (Bracci et al., 2019): nine triads, each of an animal, a lookalike and an inanimate object, framed in the category colour. None was used in training. (e) The animal bias and object bias model RDMs, which differ only in whether lookalikes are grouped with animals or with inanimate objects; dark cells mark image pairs predicted to be dissimilar. All images adapted from Bracci et al. (2019) (CC BY 4.0).

After participants responded, according to the accuracy and speed of their responses, they received feedback (correct, incorrect, or too-slow) and gained points out of 20 to make the training task more engaging (Figure 2b). The feedback screen stayed on the screen for 1250 ms. Each block was followed by a 30-second resting period, after which participants pressed the space bar to continue the experiment.

### 2.4 Behavioural Analysis

Behavioural data from the training phase were analysed using R version 4.5.2 (R Core Team, 2025). Only participants who completed all three sessions were included in the analysis. To determine the reaction time (RT) below which responses could not be reliably distinguished from guessing, we pooled all trials across all participants and sessions and grouped them into 25 ms RT bins (bins with fewer than 20 trials were excluded). For each bin, a one-tailed binomial test compared the proportion of correct responses against chance level (50%). The RT cutoff was defined as the first bin at which accuracy was significantly above chance and remained so in the subsequent bin, yielding an optimal cutoff of 287.5 ms. Trials with RTs below this threshold were excluded from all further analyses.

#### 2.4.1 Pre-Post Within-session Training Comparison

To examine immediate effects of the training, we compared performance between extra before and extra after blocks (see Section 2.3) for each category (animals, lookalikes, and objects). We analysed four measures: accuracy, defined as the proportion of correct responses among answered trials (trials on which participants responded too slowly were not scored for accuracy); RT of correct responses; IES (mean RT of correct responses divided by the proportion correct among answered trials); and slowness (the proportion of trials that exceeded the time limit). These were analysed using 2 *×* 3 repeated-measures ANOVAs (block *×* category). If the block *×* category interaction was significant, pairwise comparisons were conducted using paired-samples t-tests (two-tailed) to determine which categories showed significant training effects; comparisons were not corrected for multiple testing, as each category tested a distinct, pre-specified hypothesis (H1). Outliers were identified by calculating, for each participant, a Z-score of their mean score within each block *×* category combination (comparing each participant’s mean to the distribution of all participants’ means for that specific block and category). Participants with (*|Z| >* 3) in any combination were excluded entirely from the corresponding analyses.

#### 2.4.2 Longitudinal Across-session Comparison

We analysed the same four measures across all three sessions for each category using only extra before blocks. The extra after blocks were excluded from the longitudinal analysis since they immediately followed training and would reflect practice effects rather than consolidation. For the longitudinal analysis, we conducted 3 *×* 3 repeated-measures ANOVAs (session *×* category), followed by paired t-tests (two-tailed) with Bonferroni correction to identify whether training effects were significant across sessions for each category. In addition, we tested whether training reduced the specific difficulty associated with categorising lookalikes, over and above any general improvement in task performance. To do this, we calculated a “lookalike processing cost” for each participant and session, defined as the IES for lookalikes minus the IES for objects. Because IES combines both speed and accuracy into a single score (with lower values indicating better performance), this difference score reflects how much less efficiently lookalikes were categorised relative to inanimate objects, independent of a participant’s general improvement over sessions. A processing cost of zero would indicate that lookalikes were categorised just as efficiently as inanimate objects, whereas a positive value indicates a residual cost specific to lookalikes. We then compared this cost between the first and third sessions using a one-tailed paired t-test, predicting that training would shrink the gap between lookalikes and objects if it succeeded in reducing the animal bias specifically (rather than just improving performance across the board). Outliers were identified by applying a Z-score criterion (*|Z| >* 3), computed separately within each session *×* category combination. Participants exceeding this criterion for any relevant session *×* category combination were excluded from the corresponding analyses, including the lookalike processing cost comparison, for which a participant was excluded if they were an outlier in the lookalike or object IES for either session. Significance was set at *α* = .05 for all tests.

### 2.5 Testing Phase (EEG)

To test animacy categorisation under EEG recording, we administered the same experimental paradigm used in Chen et al. (2023) (https://osf.io/d5egu/), using a stimulus set created by Bracci et al. (2019). The stimulus set consisted of 27 greyscale images, nine per category (animals, lookalikes, and inanimate objects; Figure 2d); these images were not included in the behavioural training phase. Participants were asked, “Does the image depict a living animal?” and were required to categorise lookalikes as inanimate, the same response as for inanimate objects. Note that Chen et al. (2023) additionally included an “appearance task”, in which lookalikes had to be categorised as (appearing as) animate; the present study included only the animacy task.

Each trial began with a stimulus displayed at the centre of the screen for 500 ms, followed by a 700 ms blank screen during which participants responded as accurately and as quickly as possible. A fixation cross then appeared for a random intertrial interval of 700–1100 ms (Figure 2c). Display time did not vary with response time or accuracy, and no feedback or points were given. Key-category mappings changed between each run. Each of the eight runs consisted of 108 trials (27 images presented four times each) and lasted approximately five minutes, for a total duration of about 40 minutes; each run began and ended with a five-second fixation cross.

Participants completed the task during EEG recording in a dimly lit room, on a BenQ XL2411-B screen (24 inches, 60 Hz, 1920 *×* 1080 pixel resolution). Stimuli were presented at a size of 324 *×* 324 pixels, corresponding to approximately 8.5*^◦^* of visual angle, at a viewing distance of approximately 60 cm, consistent with Chen et al. (2023). A chin/forehead rest, positioned approximately 60 cm from the monitor, was used to minimise head movements and maintain a stable viewing distance.

#### 2.5.1 EEG Recording

We used the same setup as Chen et al. (2023). EEG was recorded using the BioSemi ActiveTwo system with 128 Ag/AgCl (silver-silver chloride) active electrodes (BioSemi, Amsterdam, Netherlands), sampled at 1024 Hz. The electrodes were placed on an elastic cap according to the International 10–20 EEG system. Cap size was determined individually for each participant based on head circumference, measured with a tape measure from just above the eyebrows around to the most prominent point at the back of the head. Participants were then fitted with a small medium (52–56 cm), medium (54–58 cm), or medium large (56–60 cm) cap accordingly. A photosensor was placed on the corner of the screen, which detected the transition between white (stimulus present) and black (stimulus not present) to record the onset of each stimulus.

### 2.6 EEG Preprocessing

Preprocessing followed the same pipeline described in Chen et al. (2023) and was carried out in MATLAB R2024b (The MathWorks Inc., 2024). After data collection, EEG and behavioural data recorded during the task were organised according to the Brain Imaging Data Structure guidelines (Gorgolewski et al., 2016) with the extension for EEG data (Pernet et al., 2019). Slow drift noise and power line noise were attenuated by applying a 2 Hz high-pass filter and a band-stop filter at 50 Hz and its harmonics (100 and 150 Hz), respectively, through the FieldTrip toolbox (Oostenveld et al., 2011). Electrode B7 was excluded because of a malfunction of the amplifier contact point for this electrode that was observed across datasets recorded with this system. Its signal was replaced by a weighted average of the neighbouring channels using FieldTrip before re-referencing. No other channels were interpolated and no epochs were rejected. Artefacts were removed with independent component analysis (see below). Traces were demeaned per run (baseline correction), which subtracts each electrode’s mean voltage across the run from its own signal, and referenced to the average of all 128 channels, which subtracts the across-electrode mean voltage at each time point from every electrode’s signal. The data were then downsampled from 1024 Hz to 250 Hz to reduce computational load without losing important signal information (Grootswagers et al., 2017).

To remove artefacts caused by eye and muscle movements, heartbeats, and channel noise, we conducted independent component analysis (ICA) using EEGLAB (Delorme & Makeig, 2004). Components were labelled using ICLabel and removed if they were classified as muscle (probability 0.6–1), eye (probability 0.4–1), heart (probability 0.6–1), or channel noise (probability 0.6–1) artefacts; components classified as brain activity, line noise, or other were retained. Following this component removal, the epoch window was set from *−*200 (before each stimulus) to 500 ms (after each stimulus), resulting in the segmentation of preprocessed data into 700 ms epochs per trial.

### 2.7 Multivariate Pattern Analysis

#### 2.7.1 Category-Level Decoding

To explore whether the neural activity patterns encode category-level discrimination, MVPA on EEG data was performed using the CosmoMVPA toolbox (Oosterhof et al., 2016). As for preprocessing, the decoding script in MATLAB was based upon the code provided by Chen et al. (2023).

We conducted condition-wise decoding for three category comparisons: animate vs. inanimate, lookalike vs. inanimate, and lookalike vs. animate. In each comparison, all trials belonging to a category were pooled under a single label, so that the classifier discriminated between two categories rather than between individual images. No trial averaging or pseudotrial construction was performed; each individual trial served as a single, independent observation for classifier training and testing. We carried out a time-resolved searchlight analysis across the epoch (*−*200 to 500 ms) using a linear discriminant analysis (LDA) classifier with leave-one-run-out cross-validation for each comparison. The classifier was trained on 504 trials (seven runs *×* 72 trials per comparison) and tested on the 72 trials of the held-out run, with this procedure repeated until each of the eight runs had served as the test set once. The temporal searchlight used multisensor patterns from all 128 sensors within a 16 ms window centred on each timepoint (2 samples on either side, given the 250 Hz sampling rate), moving forward by 4 ms (one sample) at each step across the epoch. Sensor activity across all timepoints within each window was concatenated into a single 640-dimensional feature vector per trial (128 sensors *×* 5 timepoints), without averaging across time.

We performed one-sample t-tests to test whether decoding accuracy differed from the 50% chance level for pairwise decoding. Following Chen et al. (2023), we conducted 100 permutations per participant by shuffling category labels, followed by group-level threshold-free cluster enhancement (TFCE) correction for multiple comparisons with 1,000 iterations (two-tailed, *z >* 1.96, *p* < .05).

#### 2.7.2 Three-Way Triad Decoding

In addition, we conducted a three-way triad decoding to determine the ability of the classifier to differentiate between all three categories simultaneously. We conducted the same temporal searchlight approach as in pairwise decoding. However, this time the LDA classifier was trained to perform three-way discrimination between animate, lookalike, and inanimate categories. First, we performed a leave-one-run-out cross-validation, in which the classifier was trained on data from seven runs and tested on the eighth. Because all runs contain the same 27 images, this analysis assessed whether the classifier could discriminate between the three categories in principle, without testing whether this discrimination generalised to novel, unseen exemplars.

To address this limitation, we subsequently conducted a leave-one-triad-out validation, where the classifier was trained on 8 triads (24 images) and tested on the 9th, held-out triad (the remaining 3 images not included in training). This procedure was carried out 9 times until each triad was used as a test set once. Leave-one-triad-out validation enabled the assessment of whether category representations generalise to novel exemplars. For both types of cross-validation, 3 *×* 3 confusion matrices were generated at each timepoint based on classification results, showing the frequency of each category’s correct classifications or misclassifications as a different category (e.g., how often lookalikes were classified as animals or objects).

Statistical significance was assessed at each time point by testing class-wise decoding accuracies against the chance level of 1/3 (33.33%), using a sign-flipping permutation procedure with TFCE correction for multiple comparisons across time points (1,000 iterations, two-tailed, *|z| >* 1.96, *p* < .05). In addition, mean decoding accuracies across the entire epoch, including the prestimulus baseline, are reported descriptively.

### 2.8 Representational Similarity Analysis

Complementary to the category-level decoding, we conducted an LDA-based RSA to assess how neural representations of each image differ from or are similar to the other images over time. Using the same temporal searchlight approach, we trained a separate binary LDA classifier for each of the 351 possible pairs of the 27 images to discriminate between that specific pair. Each classifier was trained and tested using leave-one-run-out cross-validation: trained on data from 7 runs and tested on the 8th, repeated until each run had served as the test set once. This was carried out independently for every image pair, for each participant, and at each time point, resulting in a 27 *×* 27 *×* 176 representational dissimilarity matrix (RDM) per participant, with each cell reflecting the classifier’s ability to discriminate that specific pair of images at that time point.

Two theoretical representational models were constructed to test whether the neural representation of lookalike objects was organised according to their animal-like visual appearance or their true inanimate identity, and whether this organisation differed following training. The animal bias and object bias models were defined as 27 *×* 27 binary RDMs, in which each cell specified the predicted relationship between a pair of images. Image pairs assigned to the same model-defined category were given a dissimilarity value of 0, whereas pairs assigned to different categories were given a value of 1.

The two models were identical in their treatment of animals and inanimate objects but differed in how they represented lookalikes. In the animal bias model, lookalikes were grouped with animals, reflecting a representational organisation based on their animal-like appearance. In the object bias model, lookalikes were instead grouped with inanimate objects, reflecting an organisation based on their identity as inanimate objects. Thus, the two model RDMs captured competing hypotheses about the position of lookalikes within neural representational space.

At each EEG time point, each participant’s neural RDM was correlated with each theoretical model using Pearson correlation. The resulting time-resolved model correlations quantified the extent to which the observed neural representational geometry corresponded to the animal bias or object bias hypothesis. Comparing these correlations across time and between groups allowed us to assess whether training was associated with a shift from an animal bias toward an object bias when representing lookalikes. Significance of the model correlations, and of the difference between them, was assessed with permutation tests with TFCE correction for multiple comparisons across time points (10,000 iterations, two-tailed, *|z| >* 1.96, *p* < .05). The mean model-correlation difference over 0–500 ms was compared between groups with an independent-samples t-test. Whether this difference changed with training was tested at each time point with a between-groups permutation test with TFCE correction (10,000 iterations, two-tailed, *|z| >* 1.96). In addition, each group’s mean difference over 0–500 ms was tested against zero with a one-sample t-test.

### 2.9 Cross-Study Comparison: Analysing Training Effect

To assess the effects of training on neural representations, we compared decoding results of the animacy task between two studies: Chen et al. (2023), which did not include a training phase, and the present study. Chen et al. (2023) had two tasks: animacy and animal appearance tasks. Only the data from the animacy task was included in the comparison since participants in the present study also performed only the animacy task. The exact same analysis pipeline of Chen et al. (2023) was applied to the present data to ensure the observed differences were due to training instead of methodological differences, adjusting only the leave-one-run-out cross-validation to account for the differing number of runs in each dataset (7 runs for the untrained group versus 8 runs for the trained group).

We defined animal bias as the difference between lookalike-inanimate and lookalike-animate decoding accuracies (see Figure 5). We computed this difference for both studies at each timepoint (176 timepoints spaced 4 ms apart, from *−*200 to 500 ms), and all the values across the 0 to 500 ms epoch were averaged. A difference between lookalike-inanimate and lookalike-animate decoding accuracies that is equal to zero means the lookalikes were classified with no preference toward either category. A negative value would mean that lookalikes were easier to distinguish from animals than from inanimate objects for the classifier, whereas a positive value would indicate that it was harder for the classifier to differentiate lookalikes from animates than inanimate objects, reflecting the animal bias. Statistical significance was assessed separately for each study using TFCE-corrected permutation tests with 1,000 iterations, applied to the per-participant difference between lookalike-inanimate and lookalike-animate decoding accuracies, to test whether this difference was significantly different from zero at the group level (two-tailed, *|z| >* 1.96). The null distribution was built from the same label-shuffling permutations used in the category-level decoding analysis. In addition, each participant’s mean animal bias over 0–500 ms was tested against zero with a one-sample t-test in each group.

Then, to test whether the animal bias differed between groups (with versus without training), the animal bias time courses (0–500 ms) of the two groups were compared using a non-parametric, between-subjects permutation test with TFCE correction and 10,000 iterations. Group labels (untrained vs. trained) were randomly shuffled across the pooled sample of 60 participants at each iteration to build the null distribution. The trained group was compared with the untrained group (trained *−* untrained), and a one-tailed threshold of *z* = *−*1.645 was applied (*p* < .05). In addition, each participant’s mean animal bias over 0–500 ms was compared between groups using a one-tailed permutation test with 100,000 iterations. For both tests, the null hypothesis stated that there was no difference in animal bias between the training and the no-training group, whereas the alternative hypothesis predicted that the animal bias would be reduced in the trained group compared to the untrained group. The random number generator was fixed (seed 1) for reproducibility.

## 3 Results

### 3.1 Behavioural Training

#### 3.1.1 Pre-Post Training Comparison

We tested in each session whether inverse efficiency scores (IES) differed between the first (extra before) and last (extra after) block (Figure 3a–c). Reaction time, slowness and accuracy were analysed in the same way and are reported in Supplementary Section S1.1 (Figures S1a–c, S2a–c and S3a–c). Before conducting the analysis, one participant was excluded from all three sessions, another participant was excluded only from Session 1, and an additional participant was excluded from Sessions 2 and 3 based on the Z-score criterion. For each session, we ran a 2 (block) *×* 3 (category) repeated-measures ANOVA. Because the question of interest was whether the training effect differed between categories, pairwise t-tests were only performed in sessions where the block *×* category interaction was significant.

**Figure 3.**
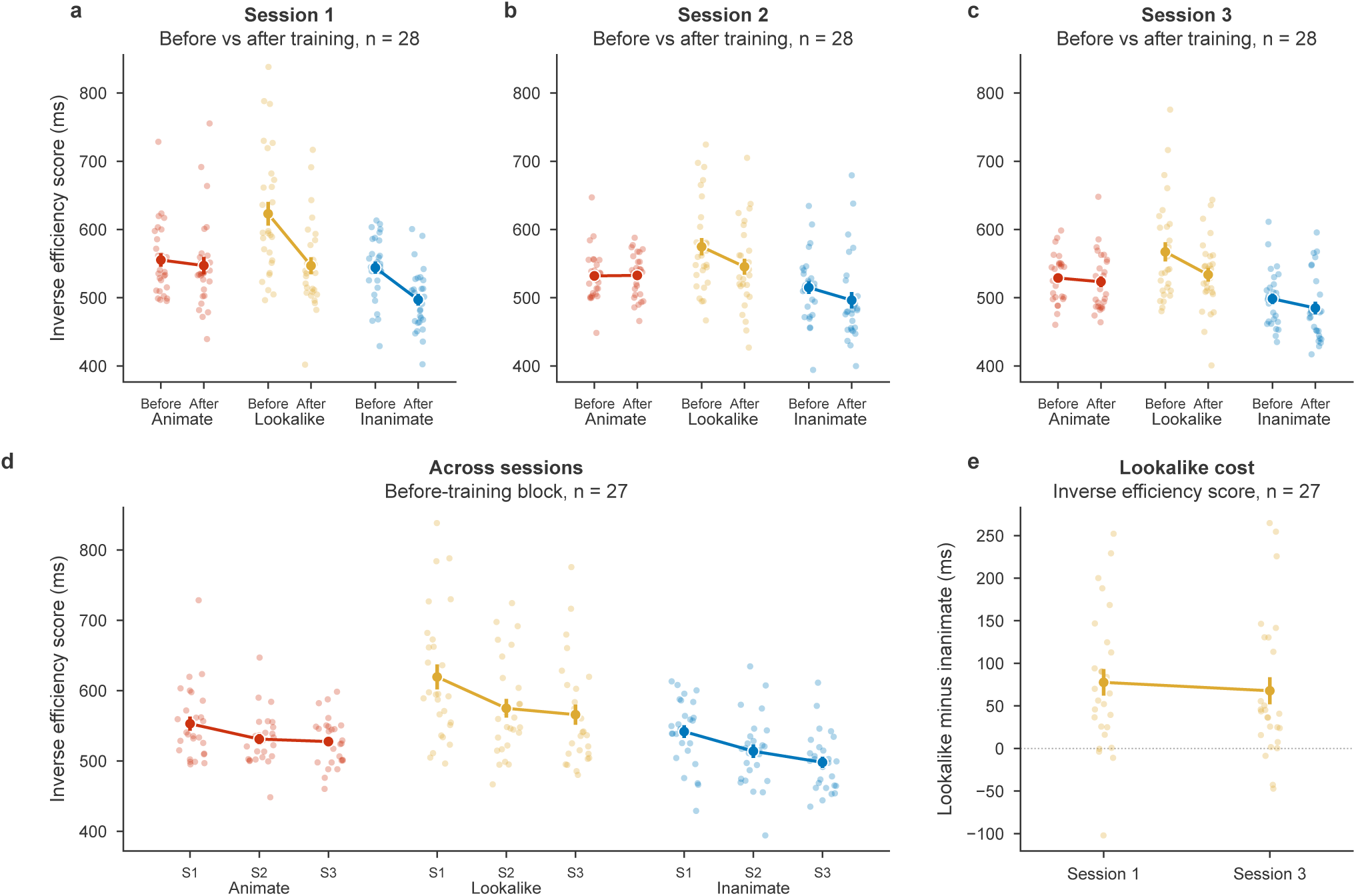
Inverse efficiency scores (IES) during behavioural training. IES is the mean reaction time of correct responses divided by the proportion of correct responses among answered trials; lower values indicate more efficient categorisation. (a–c) IES in the extra before and extra after blocks (fixed display time of 600 ms) of Sessions 1–3, for animals (animate), lookalikes and objects (inanimate). (d) IES in the extra before block of each session. (e) Lookalike processing cost, the IES for lookalikes minus the IES for objects, in Sessions 1 and 3. Dots show individual participants; filled circles and error bars show the group mean and standard error of the mean. Participants excluded by the Z-score criterion are not shown, and each panel title gives the number of participants included.

In the first session, the main effects of block [*F* (1, 27) = 42.85, *p* < .001, ƞ^2^_G_ = .109] and category [*F* (2, 54) = 17.39, *p* < .001, ƞ^2^_G_ = .151], as well as the block *×* category interaction [*F* (2, 54) = 5.68, *p* = .006, ƞ^2^_G_ = .047], were significant. Pairwise t-tests showed that IES significantly improved from extra before to extra after for lookalikes (*t*(27) = 4.54, *p* < .001, 95% CI [.042, .110]) and objects (*t*(27) = 5.23, *p* < .001, 95% CI [.029, .066]), but not for animals (*t*(27) = .61, *p* = .549, 95% CI [*−*.020, .036]).

In the second session, the main effects of block [*F* (1, 27) = 6.20, *p* = .019, ƞ^2^_G_ = .022] and category [*F* (2, 54) = 13.51, *p* < .001, ƞ^2^_G_ = .152] were significant, but the block *×* category interaction was not [*F* (1.39, 37.6) = 2.44, *p* = .117]. The same pattern was found in the third session, with significant main effects of block [*F* (1, 27) = 7.22, *p* = .012, ƞ^2^_G_ = .030] and category [*F* (2, 54) = 19.86, *p* < .001, ƞ^2^_G_ = .082] and no significant interaction [*F* (2, 54) = 1.85, *p* = .166]. In these two sessions, IES thus improved from before to after training, without reliable evidence that the size of this improvement differed between categories.

Overall, IES improved from before to after training in each session, as shown by a significant main effect of block in all three sessions. The block *×* category interaction was significant only in Session 1, where animals showed no improvement while lookalikes and objects both did. In Sessions 2 and 3, there was no reliable evidence that the improvement differed between categories. We therefore found no strong indications that training effects differed between categories, and no indication that the improvement was specific to lookalikes.

#### 3.1.2 Longitudinal Analysis

In this analysis, we test for longer-term changes across the three training sessions. The corresponding analyses of reaction time, slowness and accuracy are reported in Supplementary Section S1.2 (Figures S1d, S2d and S3d). Three participants were excluded from the analysis based on the Z-score criterion. A 3 (session) *×* 3 (category) repeated-measures ANOVA showed significant main effects of session [*F* (2, 52) = 14.74, *p* < .001, ƞ^2^_G_ = .029] and category [*F* (2, 52) = 46.54, *p* < .001, ƞ^2^_G_ = .082]. The session *×* category interaction was not significant [*F* (2.75, 71.43) = .60, *p* = .604], so there was no reliable evidence that the change in IES across sessions differed between categories (Figure 3d). Given the absence of an interaction, we did not run pairwise comparisons between sessions for individual categories. Additionally, we tested whether training decreased the perceptual cost of lookalike processing, defined as the IES difference between lookalikes and objects, from Session 1 to Session 3. This cost was numerically reduced by 10 ms, but the reduction was not statistically significant (one-tailed paired t-test, *p* = .340; Figure 3e). Together with the absence of a session *×* category interaction, this suggests that processing efficiency improved across sessions in general, without strong indications that training effects differed between categories. We therefore found no evidence that training reduced the specific processing cost of lookalikes relative to objects.

### 3.2 EEG Analyses

#### 3.2.1 Category-Level Decoding

Time-resolved pairwise decoding analysis was performed to assess whether brain activity patterns encoded information regarding animacy categories. It showed that all three pairwise comparisons, including animate versus inanimate, lookalike versus inanimate, and lookalike versus animate, reached significant above-chance decoding accuracy following stimulus onset at multiple time points, confirming that for all three pairwise comparisons the EEG signal contained category-level information (Figure 4a). Significant above-chance decoding was first observed at 80 ms for animate versus inanimate, at 76 ms for lookalike versus inanimate, and at 96 ms for lookalike versus animate (*p* < .05, TFCE-corrected).

**Figure 4.**
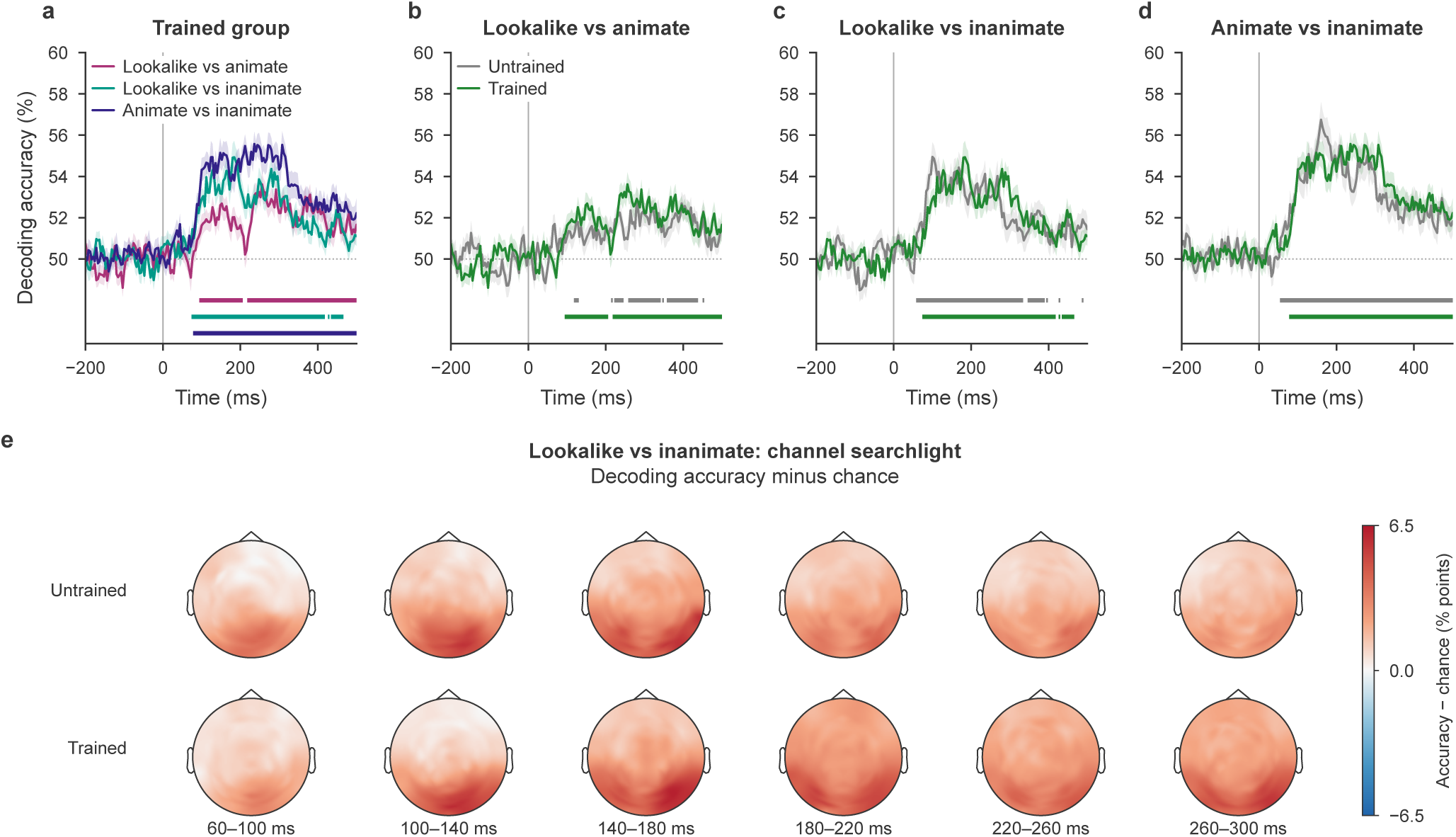
Time-resolved pairwise category decoding. (a) Trained group, all three category comparisons. (b–d) Each comparison in the untrained group (Chen et al., 2023) and the trained group: lookalike versus animate (b), lookalike versus inanimate (c) and animate versus inanimate (d). Lines show the group mean and shaded areas the standard error of the mean. Bars below the curves mark time points of significant above-chance decoding (*p* < .05, TFCE-corrected), in the colour of their curve. The dotted line marks chance (50%), and the vertical line stimulus onset. (e) Scalp distribution of lookalike versus inanimate decoding: for every channel, decoding was computed on that channel and its nine nearest neighbours (*±*8 ms), and the group mean accuracy minus chance is shown for the untrained (top) and trained (bottom) groups, in 40 ms windows from 60 to 300 ms. These maps are descriptive; no statistical test was run on them.

Among the three comparisons, animate-inanimate pairwise decoding had a peak accuracy of 55.57% at 236 ms, and it remained significant until the end of the epoch (106 out of 176 time points), making it the most sustained decoding accuracy. Lookalike-inanimate pairwise decoding reached a peak accuracy of 54.93% at 184 ms, and its significance remained until 464 ms (95 out of 176 time points). In general, lookalike-animate pairwise decoding showed the weakest decoding with a peak at 53.62% at 256 ms (99 out of 176 time points). It remained significantly above-chance until the end of the epoch. This weaker decoding suggests a lower neural separability between animals and lookalikes.

Figure 4b–d explicitly compares the decoding in each of the three comparisons between the trained group (present study) and the untrained group (Chen et al., 2023). The similarities are striking. Across all three condition pairs–animate-inanimate, lookalike-inanimate, and lookalike-animate–the trained group displayed highly similar decoding trajectories with the untrained group throughout the post-stimulus window.

### Training Effects on Animal Bias

As explained before, we quantified the animal bias from the difference between lookalike-inanimate and lookalike-animate. This difference was positive on average in the post-stimulus window in both the untrained (*t*(29) = 5.13*, p* < .001) and trained samples (*t*(29) = 3.74*, p* < .001, Figure 5b), suggesting that the neural patterns associated with lookalikes were less distinct from animals than from inanimates. Statistical significance was also established using within-subjects permutation tests with TFCE correction, applied directly to the difference timecourse against zero, which confirmed the presence of a significant animal bias in both samples.

**Figure 5.**
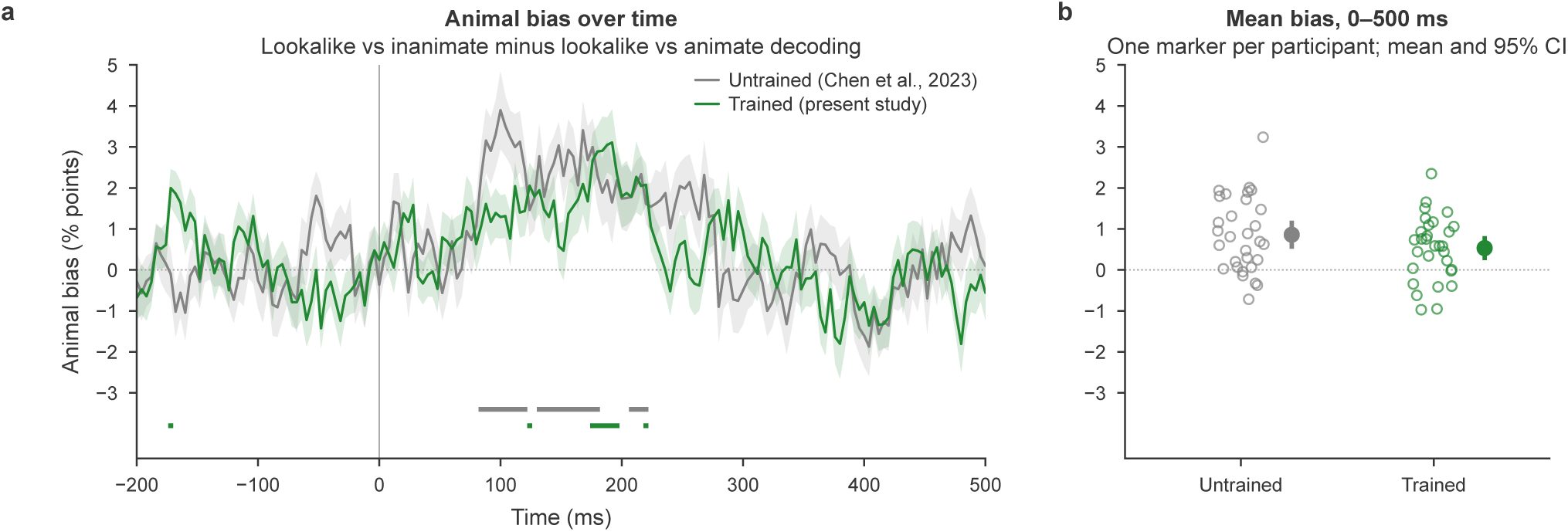
Animal bias in the untrained and trained groups. (a) Animal bias over time: lookalike-versus-inanimate minus lookalike-versus-animate decoding accuracy. Positive values mean that lookalikes are harder to distinguish from animals than from inanimate objects. Lines show the group mean and shaded areas the standard error of the mean. Bars below the curves mark time points where the bias differs from zero within each group (*p* < .05, TFCE-corrected); no time point differed between the groups (between-groups test over 0–500 ms, TFCE-corrected). (b) Each participant’s mean bias over 0–500 ms (open circles), with the group mean and 95% confidence interval (filled circle and error bars).

For the group without training (Chen et al., 2023), the temporal window where animal bias reached statistical significance was between 84 and 220 ms after the presentation of the stimulus (27 out of 176 time points, *p* < .05, TFCE-corrected) with a peak difference of 3.90 percentage points at 100 ms and a mean difference between lookalike-inanimate and lookalike-animate decoding accuracies of .0086, indicating a 0.86-percentage-point difference across the post-stimulus window. For the trained group, the animal bias was statistically significant at 9 out of 176 time points (*p* < .05, TFCE-corrected). Most of these fell within a narrower temporal window between 176 and 196 ms post-stimulus onset (6 time points), with a peak difference of 3.11 percentage points at 192 ms. Two further isolated time points were significant at 124 and 220 ms. In addition, one isolated time point reached significance before stimulus onset (*−*172 ms). Because no stimulus information is available at this latency, it most likely reflects noise rather than a genuine effect. The difference between lookalike-inanimate and lookalike-animate decoding in this trained group was .0053, indicating a 0.53-percentage-point difference across the post-stimulus window. A TFCE-corrected Monte Carlo permutation test with 10,000 iterations demonstrated that there was no statistically significant difference between the two groups at any time point between 0 and 500 ms (one-tailed; smallest *p* = .201 at 108 ms; two-tailed: *p* = .402). Likewise, when the animal bias was averaged over the 0–500 ms window for each participant, it did not differ significantly between the two groups (one-tailed permutation test, 100,000 iterations, *p* = .071). In summary, both groups showed a significant animal bias, and this bias was not significantly different between the two groups.

#### 3.2.2 Three-Way Triad Decoding

A three-way triad decoding analysis was performed to assess whether the classifier could differentiate between animate, lookalike, and inanimate categories simultaneously in the group with training. According to the results of leave-one-run-out cross-validation, the decoding accuracies of all three categories exceeded chance level at multiple time points following stimulus onset with an overall mean decoding accuracy of 35.14%. The decoding accuracy was 35.22% (significant at 105 out of 176 time points) for animates with a peak at 38.37% at 248 ms, 35.75% (significant at 100 out of 176 time points) for inanimates with a peak at 39.94% at 188 ms, and 34.45% (significant at 70 out of 176 time points) for lookalikes with a peak at 36.56% at 276 ms (Figure 6a). Descriptively, inanimate objects reached the highest numerical peak decoding accuracy, followed by animates and lookalikes. This is what one expects to find in the case of an animal bias, which would cluster animate objects and lookalikes and make inanimate objects the outlier category that is most easily discriminated from the two other categories.

**Figure 6.**
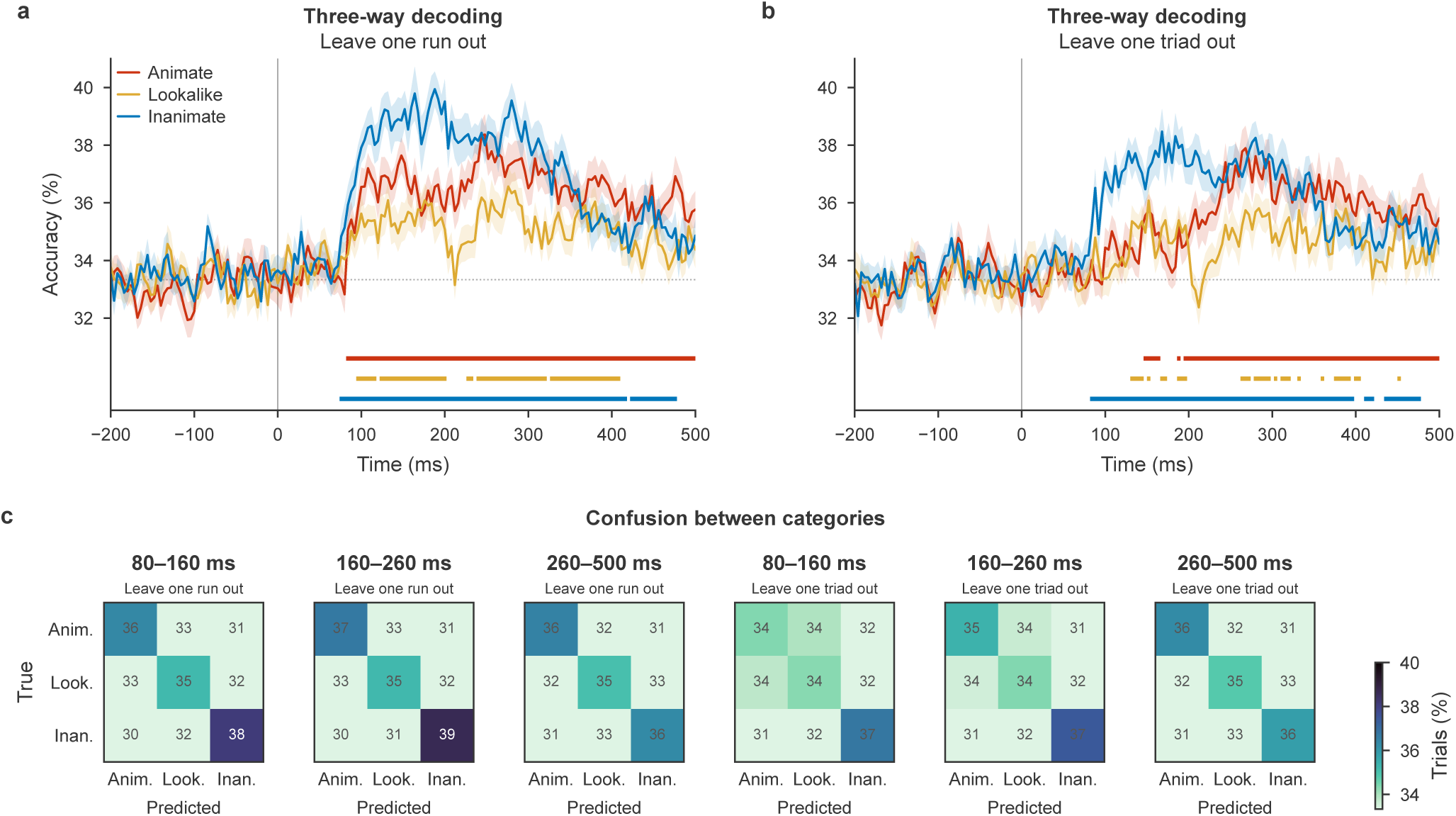
Three-way category decoding in the trained group. (a) Class-wise decoding accuracy with leave-one-run-out cross-validation, in which the classifier was trained on seven runs and tested on the held-out run. (b) Leave-one-triad-out cross-validation, in which the classifier was trained on eight triads and tested on the held-out triad, a test of generalisation to unseen exemplars. Lines show the group mean and shaded areas the standard error of the mean; bars below the curves mark time points of significant above-chance decoding (*p* < .05, TFCE-corrected); the dotted line marks chance (33.3%). (c) Group confusion matrices in three time windows (80–160, 160–260 and 260–500 ms) for both cross-validation schemes: rows are the true category, columns the predicted category, and cells the percentage of trials.

This leave-one-run-out analysis demonstrated that EEG activity contained information sufficient to discriminate among the three stimulus categories. However, because the same 27 images appeared in both the training and test sets across runs, successful decoding could partly reflect image-specific information rather than representations that generalised across exemplars. We therefore performed a leave-one-triad-out cross-validation analysis, in which the classifier was tested on exemplars that were excluded entirely from classifier training. This provided a more stringent test of whether category-discriminative information generalised to unseen exemplars.

Using leave-one-triad-out cross-validation, mean class-wise decoding accuracy was 34.73% for animates, with significant above-chance decoding at 83 of 176 time points and a peak accuracy of 37.89% at 268 ms. For inanimates, mean decoding accuracy was 35.26%, with 93 significant time points and a peak of 38.47% at 168 ms. Lookalikes showed a mean decoding accuracy of 34.09%, with 32 significant time points and a peak of 36.08% at 152 ms (Figure 6b). Overall mean decoding accuracy was 34.69%, compared with a nominal chance level of 33.33%. The presence of above-chance decoding for exemplars excluded from classifier training indicates that the discriminative neural information was not restricted to the specific images used to train the classifier and showed some generalisation across exemplars. Again, the three-way decoding is descriptively highest for inanimate objects, more specifically in the earlier time points until around 200 ms.

#### 3.2.3 Representational Similarity Analysis

Previous decoding analyses demonstrated that EEG activity contained information sufficient to discriminate between the animacy categories. However, while decoding establishes whether neural signals can distinguish between categories, it does not reveal how these stimuli are organised relative to one another in representational space. Therefore, an LDA-based RSA was conducted to map the overall similarity structure of all the 27 images over time. In particular, we investigated whether neural representations were organised in line with an animal bias or an object bias. To address this question, EEG RDMs were correlated with two theoretical models, the animal bias model and the object bias model.

In Chen et al. (2023), the association of the animal bias model with RDMs was strong and early, with significant correlations first observed at 68 ms and a peak at *r* = .12 at 176 ms (74 out of 176 time points, *p* < .05, TFCE-corrected). The object bias model demonstrated a weaker and more delayed correlation compared to the animal bias model, with significant correlations first observed at 152 ms and a peak at *r* = .06 at 168 ms (18 out of 176 time points, *p* < .05, TFCE-corrected) (Figure 7c). A within-group one-sample permutation test with TFCE correction and 10,000 iterations was conducted to assess whether the object bias model and the animal bias model significantly differ from each other in Chen et al. (2023). It showed that the difference between the two models during the 144–232 ms window was statistically significant at several time points (12 out of 176 time points, *p* < .05, TFCE-corrected), showing a benefit for the animal bias model.

**Figure 7.**
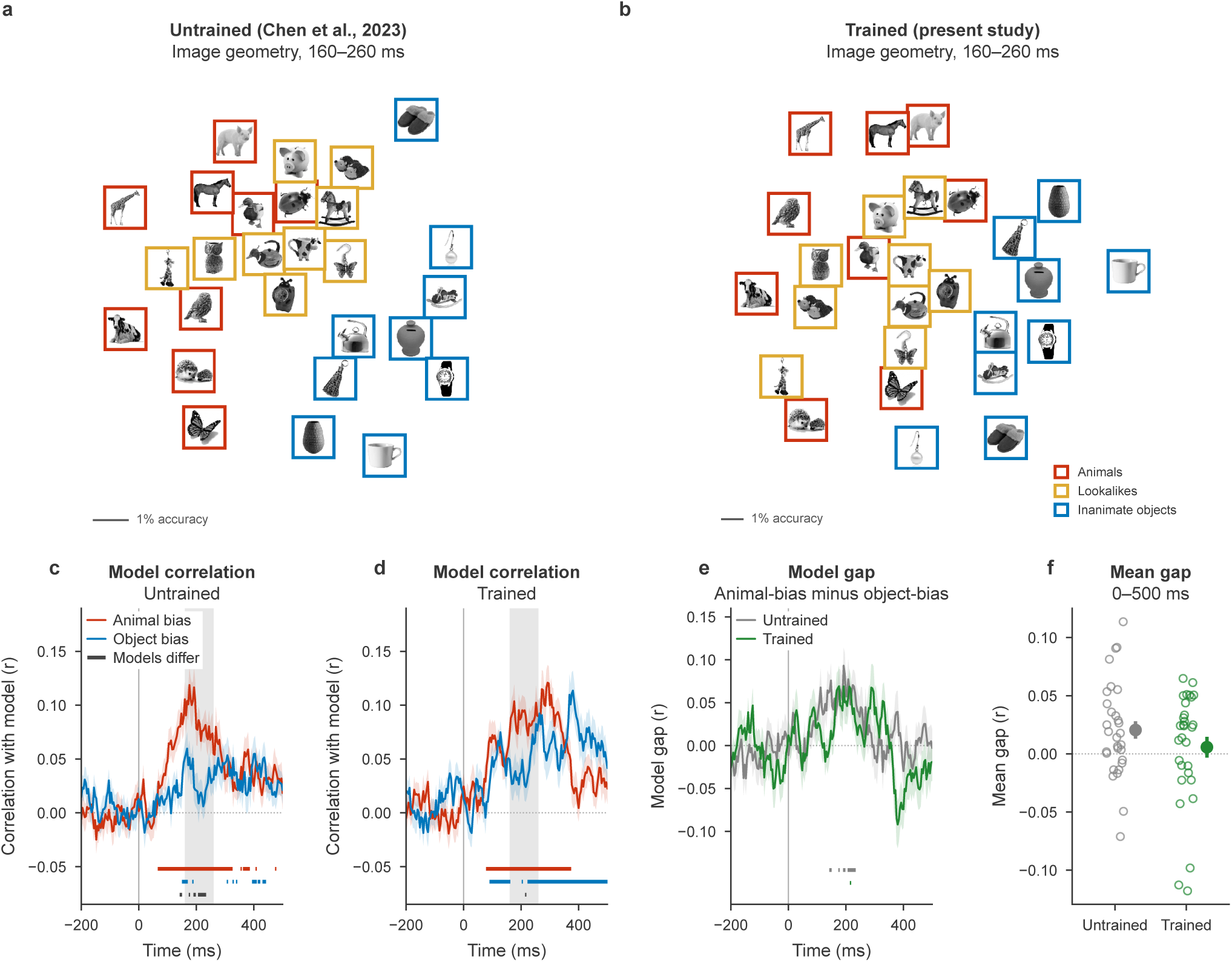
Representational geometry of the 27 images and model RSA. (a, b) Metric multidimensional scaling of the group-mean neural RDM (pairwise image decoding accuracy above chance) at 160–260 ms, for the untrained group (Chen et al., 2023) (a) and the trained group (b). Each thumbnail sits at its image’s position, nudged aside only where thumbnails would overlap and then joined to its true position by a thin line; the frame colour gives the category (red: animals; yellow: lookalikes; blue: inanimate objects). The trained solution is rotated onto the untrained one, and the scale bar marks one percentage point of decoding accuracy. (c, d) Correlation of each participant’s neural RDM with the animal bias and object bias models over time, in the untrained (c) and trained (d) groups. Coloured bars mark significant correlations with each model and the dark bar marks where the two model correlations differ; the grey band marks the time window shown in a and b. (e) Difference between the animal bias and object bias model correlations in both groups. Bars in each group’s colour mark where that group’s difference is significant, and dark bars would mark a difference between the groups. (f) Each participant’s mean difference over 0–500 ms, with the group mean and standard error of the mean. In c–e, lines show the group mean and shaded areas the standard error of the mean, and all bars are *p* < .05, TFCE-corrected.

In the present study, significant correlations for the animal bias model were first observed at 80 ms with a peak at *r* = .12 at 292 ms (74 out of 176 time points, *p* < .05, TFCE-corrected). In addition, significant correlations for the object bias model were first observed at 92 ms and a peak at *r* = .11 at 380 ms (89 out of 176 time points, *p* < .05, TFCE-corrected) (Figure 7d). A within-group one-sample permutation test with TFCE correction and 10,000 iterations revealed that the two models differed significantly only at a single time point, at 216 ms, showing a brief benefit for the animal bias model. The animal bias model remained comparable across the groups with and without training. However, the correlation between the object bias model and RDMs included a higher peak (with training: *r* = .11 versus without training: *r* = .06) and a greater number of significant time points (with training: 89 out of 176 time points versus without training: 18 out of 176 time points) in the trained group relative to the untrained group.

A between-groups permutation test with TFCE correction and 10,000 iterations was performed to investigate whether the difference between the animal bias model and the object bias model correlations changed between the group with training and without training. This revealed no significant difference between the two groups at any time point across the epoch. The largest difference occurred at 380 ms but did not reach the two-tailed threshold (*z* = 1.88) (Figure 7e). To investigate whether the overall magnitude of the animal bias changed following training, the difference between the animal bias and object bias model correlations was averaged across the 0–500 ms post-stimulus window (Figure 7f). The untrained group exhibited a mean difference of .021, which was significantly different from zero (*t*(29) = 2.74, *p* = .010), whereas the trained group exhibited a mean difference of .006, which was not (*t*(29) = 0.65, *p* = .522). The difference between groups was not significant (independent-samples t-test, *t*(58) = 1.27, *p* = .210). It should be noted that the small mean difference in the trained group reflects two opposing effects across the epoch (Figure 7e). In the early time window, the difference was positive, indicating an animal bias, whereas around 350–450 ms it became negative, indicating an object bias. These two effects partly canceled each other out when the difference was averaged across the 0–500 ms window.

## 4 Discussion

The present study investigated whether short-term, feedback-based categorisation training could attenuate the visual system’s animal bias. Behaviourally, while participants demonstrated within-session learning and general task improvement, the processing cost associated with lookalikes showed little longterm reduction. This behavioural stability was also reflected at the neural level. Time-resolved decoding and representational similarity analysis revealed no statistically reliable attenuation of the animal bias neural signature. In RSA, the object bias model was descriptively more strongly associated with the neural data after training, but this did not result in a significant difference between groups. A significant animal bias nevertheless persisted in the trained group despite a training regime designed to counteract it (controversial training).

Beyond being a primary metric of learning, behavioural efficiency of animacy categorisation has also been shown to couple with the distinguishability of neural decoding patterns (T. A. Carlson et al., 2014; Ritchie et al., 2015). The analysis of the training data showed that participants became more efficient at the categorisation task within each session. In the first session, inverse efficiency scores improved for lookalikes and inanimate objects, while performance for animals remained stable (Figure 3a). In the second and third sessions, efficiency still improved from before to after training, but there was no reliable evidence that this improvement differed between categories (Figure 3b–c).

The first-session pattern broadly aligns with models of visual perceptual learning. Because real animals may elicit less category conflict than lookalikes, baseline categorisation performance for these stimuli may leave less room for improvement. However, lookalikes act as “boundary-adjacent exemplars”, images that sit on the edge between two categories, and, as O’Bryan et al. (2024) demonstrated, active category learning does not improve perception uniformly across categories. Instead, improvements can be particularly pronounced for these boundary-adjacent items. Nevertheless, because inanimate objects improved as well, this improvement cannot be attributed specifically to lookalikes.

This improvement in inanimate objects may reflect broader mechanisms of dimensional modulation. According to Folstein et al. (2015), learning category boundaries reorganises visual representations by increasing separability along task-relevant dimensions, a shift in the features participants relied on. Active training to distinguish animates from inanimates may result in changes consistent with a transformation of the representational space to increase the discriminability of diagnostic inanimate features such as texture. Such a shift could contribute to more efficient categorisation of stimuli possessing these features, although the behavioural results alone cannot establish representational reorganisation.

Longitudinal results showed that categorisation efficiency improved across the three sessions in general, as indicated by a significant main effect of session. This suggests that the training procedure engaged the participants and is consistent with performance gains being retained between the weekly sessions. However, this improvement did not differ between categories.

On the other hand, longitudinal behavioural analysis also reveals a limitation. Despite this general improvement in the categorisation task, the performance gap between categorising a lookalike and an inanimate object did not significantly decrease from the first to the third session (Figure 3d–e). This suggests a limited effect of training on resolving category ambiguity, which highlights the strength of the animal bias. Because lookalikes share the shape of animals, their appearance may continue to interfere with categorisation according to their inanimate identity. Overall, the results suggest that even though we observed long-term task learning in general, we did not find a statistically reliable attenuation of the specific costs associated with lookalikes.

At the neural level, the absence of a clear training-induced reduction in the behavioural animal bias was paralleled by the persistence of the neural animal bias. Our central prediction was that, if feedback-based categorisation training could modify the representations underlying the bias, lookalikes should become neurally more distinguishable from animals and more similar to inanimate objects. However, the overall attenuation of the animal bias did not reach statistical significance in the between-group comparison. Thus, although the trained group showed descriptive changes in the predicted direction, the present results do not provide statistically reliable evidence that three sessions of categorisation training altered the neural bias elicited by animal-like appearance.

Beyond its overall magnitude, the temporal profile of the animal bias provides an opportunity to examine when these representations are expressed during visual processing. In the untrained group, the animal bias, measured as the difference between lookalike-inanimate and lookalike-animate pairwise decoding, was significant at multiple time points between 84 and 220 ms after stimulus onset. In the trained group, significant effects were observed at fewer time points, primarily between 176 and 196 ms, with additional isolated significant time points at 124 and 220 ms (Figure 5). Although these patterns differ descriptively, the absence of statistically significant between-group differences prevents us from concluding that training altered the temporal dynamics of the animal bias. Previous work has shown that early EEG decoding signals reflect the rapid emergence of visual object information (Contini et al., 2017). Furthermore, Chen et al. (2023) demonstrated through fMRI-EEG fusion that the early animal bias is associated with posterior regions of the VVS, while anterior VVS regions contribute to later processing stages. Whether training differentially affects these stages remains unresolved and could be tested in future studies combining training with fMRI-EEG fusion.

Another primary hypothesis of the current study was to assess whether categorisation training altered the balance between the animal bias model and the object bias model at the level of neural representational geometry, as indexed by RSA model correlations over time. The results reveal a descriptive pattern consistent with such a shift, though the between-group evidence does not reliably confirm it. In the untrained baseline sample of Chen et al. (2023), the animal bias model showed a strong and early correlation with neural RDMs, while the object bias model was weaker and more fragmented. The two models differed significantly from one another at several time points between 144 and 232 ms, consistent with stronger representation of lookalikes according to appearance than categorical identity. This pattern aligns with the animal bias observed in the pairwise decoding results of the same study. In the trained group, the animal bias model correlation was descriptively similar to that in the untrained group.

While the animal bias model correlation was descriptively similar across groups, the object bias model showed a descriptively stronger and more sustained correlation with neural RDMs compared to the untrained group (Figure 7). The two model correlations differed significantly at only a single time point in the trained group (216 ms), compared with a longer window in the untrained baseline.

In the trained sample, the highest peak of the object bias model’s correlation occurs at 380 ms, a latency consistent with later stages of visual object processing and semantic integration (Contini et al., 2017). This interpretation is also consistent with the fMRI-EEG fusion results of Chen et al. (2023), which associated later processing with anterior regions of the VVS. Furthermore, they reported that this late-stage processing is also sensitive to task context. Only when participants actively performed an animacy task (judging biological identity), rather than an appearance task (judging visual resemblance), did the correlation with the object bias model reach levels comparable to those of the animal bias model (Chen et al., 2023).

The between-group permutation test on the difference between object bias and animal bias model correlations did not show a significant difference at any time point (Figure 7e). Therefore, although the object bias model was descriptively stronger in the late time window, the RSA findings do not provide statistically reliable evidence for a training-induced shift.

The perceptual learning literature provided theoretical grounds for expecting training-induced plasticity. O’Bryan et al. (2024) demonstrated that active category learning induces functional plasticity within early visual cortex (V1–V3). Ester et al. (2020) similarly found that category learning alters early visual processing, causing representational sharpening specifically for boundary-adjacent stimuli, potentially involving task-dependent attentional gain on diagnostic features (Byers & Serences, 2014; Van Gulick & Gauthier, 2014). However, much of this evidence concerns simple, unidimensional stimuli such as oriented bars.

Furthermore, even when training involves complex objects, plasticity has been observed when the visual system has no strongly established representation of the trained stimuli. For example, Clarke et al. (2016) showed that VVS representations reorganise after learning semantic associations of complex novel objects called fribbles. The animal bias, by contrast, involves representations of familiar animal-like visual features. The failure to find a reliable bias reduction may therefore reflect a boundary condition of short-term perceptual plasticity, one where the prior is not a simple feature or a novel object open to fast categorisation but an established representational structure that brief training may be less able to alter.

One possible explanation for this limited attenuation is that the animal bias reflects a representational structure consolidated through extensive visual experience. Duyck, Costantino et al. (2024) suggest that the human animal bias may emerge from learning the statistical regularities associated with animal appearance over a lifetime. Under this account, repeated exposure to diagnostic animal-like features may strengthen their association with neural representations of genuine animate stimuli. Lookalikes may therefore be particularly difficult to dissociate from animals because they recruit overlapping representational mechanisms, consistent with the shared computational machinery account proposed in the Introduction (H2). Such representations may become relatively resistant to short-term training, even when behavioural categorisation improves.

This account is not the only possible explanation for the persistence of the animal bias. The bias has also been discussed in relation to evolutionary pressures favouring detection of animate agents and a developmental history of exposure to animal shapes (Bracci et al., 2019; Chen et al., 2023; Duyck et al., 2024). It remains to be determined whether this resistance is primarily dictated by evolutionary factors, developmental consolidation, overlapping computational networks, insufficient training intensity, or an interaction among some or all of these. Given recent findings that the strength of the animal bias depends upon how much a lookalike object looks like an animal (Rostami et al., 2026), this resistance might also depend upon such stimulus characteristics.

A primary methodological limitation of the current study pertains to the temporal spacing and overall intensity of the categorisation training. Our protocol was distributed across three weeks, with participants completing one session per week. While this longitudinal design was valuable for examining whether learning effects persisted across sessions, the total amount of training was relatively limited. Whether longer or more intensive training, or a different distribution of sessions, would be sufficient to modify the animal bias remains an open question.

Another methodological limitation of the current investigation is the reliance on a cross-study, between-groups design to assess neural plasticity. Our use of the difference between lookalike-inanimate and lookalike-animate decoding accuracies may mitigate the influence of global differences in decoding performance between our trained group and the untrained group (Chen et al., 2023). We also used the same EEG machine and applied a closely matched analysis pipeline to minimise methodological variance and strengthen the interpretability of the findings. Nevertheless, between-group comparisons remain limited by individual differences in baseline neural representations and other potential between-study differences. Because we did not measure the initial strength of the animal bias in our trained group prior to training, we cannot determine the extent to which pre-existing differences between the groups may have influenced the results. A within-subjects design, in which the same participants undergo EEG recordings both before and after training, would address this limitation by allowing a more direct assessment of learning-induced representational changes.

Overall, the present findings suggest that the animal bias may be relatively resistant to short-term perceptual training. Despite three sessions of active feedback-based training, the behavioural processing cost of lookalike stimuli relative to inanimate objects did not significantly decrease across sessions, and neither pairwise decoding nor RSA revealed a statistically reliable attenuation of the neural bias relative to the untrained group. While within-session improvement indicated continued behavioural learning, it did not coincide with reliable evidence for a change in the neural animal bias. These findings highlight potential limits to short-term modification of established visual representations, while leaving open whether longer or more intensive training could alter them. Identifying the conditions under which the animal bias can be modified will help clarify the limits of experience-dependent plasticity in the human visual system.

## Declaration of the Use of AI

AI tools (Claude Sonnet 5 and Claude Opus 5.5, Anthropic; GPT-5 and GPT-6, OpenAI; and Grammarly) were used for copyediting, grammar and spelling checks, as well as to assist with debugging, simplifying and improving code quality and refining figure presentation; all code was reviewed and run by the authors. AI played no role in study design, data collection, initial analyses, or interpretation of the results.

## Data and Code Availability

The behavioural and EEG data supporting the findings of this study, along with the associated analysis code (PsychoPy experimental paradigm, MATLAB EEG preprocessing and MVPA/RSA pipelines, and R behavioural analysis scripts), will be made available via the KU Leuven Research Data Repository (RDR; https://rdr.kuleuven.be), in accordance with the terms of the informed consent obtained under ethical approval G-2024-8866-R2(MIN) from the Social and Societal Ethics Committee of KU Leuven.

## Author Contributions

Leylan Öykü Etkin: Methodology, Software, Investigation, Formal Analysis, Data Curation, Visualisation, Writing – Original Draft, Writing – Review & Editing. Enes Bahadır Kömürcü: Methodology, Software, Investigation, Formal Analysis, Data Curation, Visualisation, Writing – Original Draft, Writing – Review & Editing. Chiu-Yueh Chen: Resources (provision of the original experimental paradigm and EEG analysis pipeline), Methodology consultation. Hans Op de Beeck: Conceptualisation, Methodology, Supervision, Funding Acquisition, Writing – Review & Editing. Andrea Ivan Costantino: Supervision, Methodology (guidance on EEG preprocessing, MVPA, and RSA pipeline implementation), Writing – Review & Editing.

## Acknowledgements

We thank Klara Schevenels for practical and technical assistance during data acquisition and for data management.

## Supplementary Materials

## S1 Behavioural Training: Reaction Time, Slowness, and Accuracy

### S1.1 Pre-Post Training Comparison

#### Reaction Time (Figure S1a–c)

Prior to analysis, the same participant was excluded from both Session 1 and 2, while no participants were removed from Session 3 according to the Z-score criterion. The mean reaction time of correct responses in extra before and extra after blocks was compared with a 2 (block) *×* 3 (category) repeated measures ANOVA to investigate whether reaction time was influenced by the immediate effects of training. This analysis was performed for all three sessions separately. In the first session, we found a significant main effect of block [*F* (1, 28) = 77.24, *p* < .001, ƞ^2^_G_ = .248] and a significant main effect of category [*F* (2, 56) = 14.15, *p* < .001, ƞ^2^_G_ = .047]. However, the block *×* category interaction was not significant [*F* (2, 56) = 1.71, *p* = .191], indicating that the magnitude of training did not differ across image categories. As the block *×* category interaction was not significant, no pairwise comparisons were performed.

**Figure S1.**
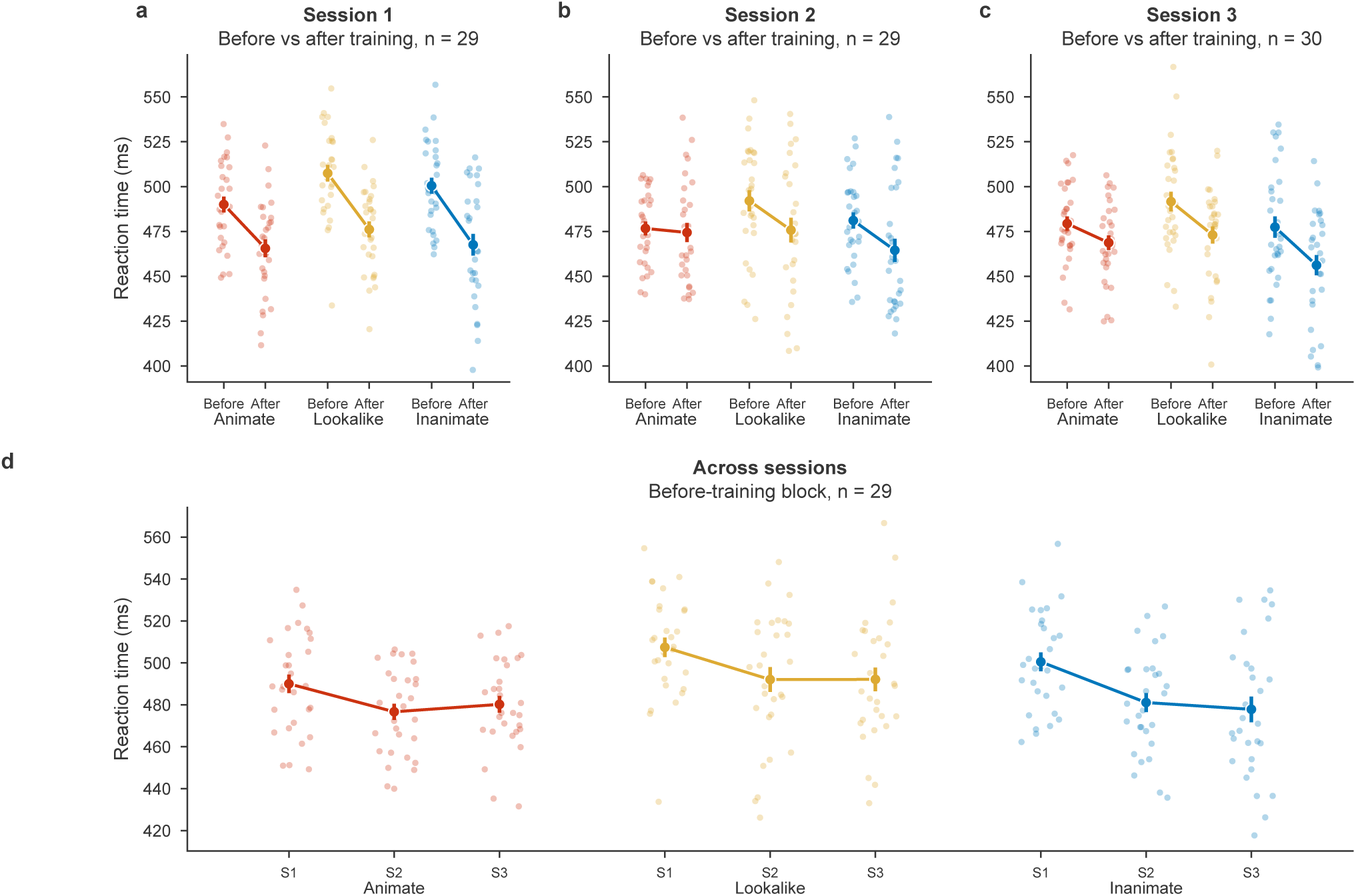
Reaction time of correct responses during behavioural training. (a–c) Extra before and extra after blocks (fixed display time of 600 ms) of Sessions 1–3, for animals (animate), lookalikes and objects (inanimate). (d) Extra before block of each session. Dots show individual participants; filled circles and error bars show the group mean and standard error of the mean. Participants excluded by the Z-score criterion are not shown, and each panel title gives the number of participants included.

In the second session, there was a significant main effect of block [*F* (1, 28) = 10.37, *p* = .003, ƞ^2^_G_ = .038], a significant main effect of category [*F* (2, 56) = 4.90, *p* = .011, ƞ^2^_G_ = .025], and a significant block *×* category interaction [*F* (2, 56) = 6.46, *p* = .003, ƞ^2^_G_ = .013]. Post hoc pairwise comparisons demonstrated that lookalikes (*t*(28) = 3.37, *p* = .002, 95% CI [.006, .026]) and objects (*t*(28) = 3.62, *p* = .001, 95% CI [.007, .026]) were significantly improved from extra before to extra after blocks regarding RT. However, there was no significant improvement for animals (*t*(28) = .56, *p* = .577, 95% CI [*−*.006, .010]). The mean improvement of RT for lookalikes and objects was 16.3 ms and 16.6 ms, respectively, whereas for animals, it was only 2.3 ms.

In the third session, a significant main effect of block [*F* (1, 29) = 14.65, *p* < .001, ƞ^2^_G_ = .088] and a significant main effect of category [*F* (2, 58) = 15.71, *p* < .001, ƞ^2^_G_ = .052] were observed. Nevertheless, there was no interaction effect between block and category [*F* (2, 58) = 2.69, *p* = .076]. As the interaction was not significant, no pairwise comparisons were performed.

#### Slowness (Figure S2a–c)

For this analysis, one participant was excluded from both Sessions 1 and 2, and another participant was excluded from both Sessions 2 and 3 based on the Z-score criterion. A 2 (block) *×* 3 (category) repeated measures ANOVA was performed to compare the proportion of trials where participants did not respond within the display time (too-slow trials) between extra before and extra after blocks. In the first session, the main effect of blocks [*F* (1, 28) = 114.72, *p* < .001, ƞ^2^_G_ = .0277], the main effect of category [*F* (2, 56) = 22.11, *p* < .001, ƞ^2^_G_ = .0123], and the interaction effect between block and category [*F* (2, 56) = 10.00, *p* < .001, ƞ^2^_G_ = .070] were significant. Pairwise t-tests revealed that the number of too-slow answers decreased significantly from extra before to extra after blocks for all categories: animals (*t*(28) = 4.17, *p* < .001, 95% CI [.018, .054]), lookalikes (*t*(28) = 8.41, *p* < .001, 95% CI [.098, .161]), and objects (*t*(28) = 5.24, *p* < .001, 95% CI [.058, .133]).

**Figure S2.**
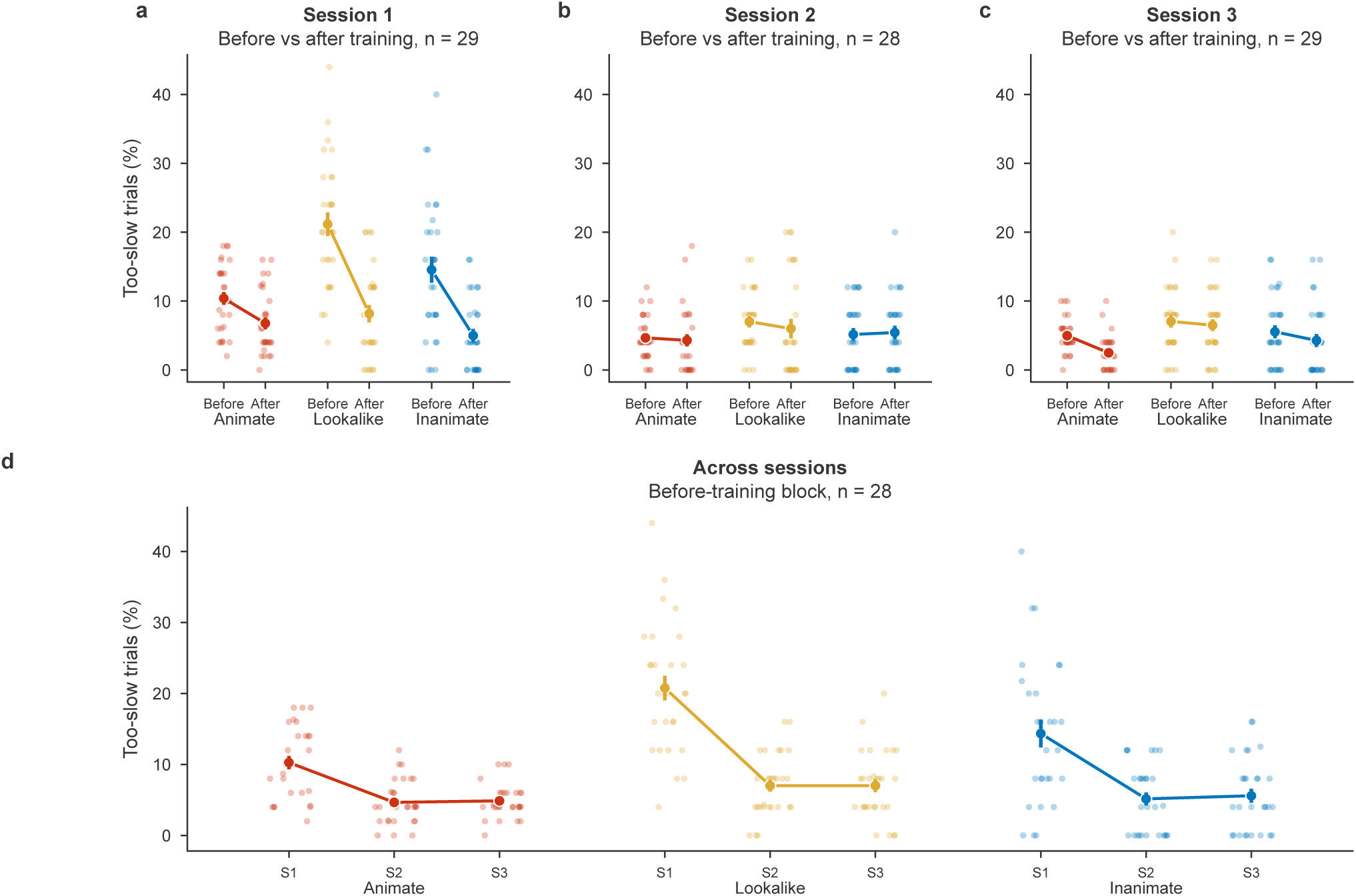
Proportion of too-slow trials (no response within the display time) during behavioural training. (a–c) Extra before and extra after blocks (fixed display time of 600 ms) of Sessions 1–3, for animals (animate), lookalikes and objects (inanimate). (d) Extra before block of each session. Dots show individual participants; filled circles and error bars show the group mean and standard error of the mean. Participants excluded by the Z-score criterion are not shown, and each panel title gives the number of participants included.

In the second session, there were no significant main effects of block [*F* (1, 27) = .31, *p* = .584] or category [*F* (1.59, 42.96) = 2.30, *p* = .123] and no significant block *×* category interaction [*F* (2, 54) = .48, *p* = .620]. Therefore, the number of answers that were marked as too-slow did not change significantly when there was training. In the third session, we observed a significant main effect of block [*F* (1, 28) = 6.41, *p* = .017, ƞ^2^_G_ = .029] and category [*F* (2, 56) = 12.41, *p* < .001, ƞ^2^_G_ = .082], whereas there was no significant interaction effect [*F* (2, 56) = 1.38, *p* = .261]. As the interaction was not significant, no pairwise comparisons were performed.

#### Accuracy (Figure S3a–c)

Prior to analysis, one participant was excluded only from Session 1, two participants were excluded from both Session 2 and Session 3, and one additional participant was excluded from only Session 3 based on the Z-score criterion. A 2 (block) *×* 3 (category) repeated measures ANOVA revealed that there was no significant main effect of block in any session for accuracy (Session 1: *F* (1, 28) = 2.01, *p* = .168; Session 2: *F* (1, 27) = .034, *p* = .856; Session 3: *F* (1, 26) = .045, *p* = .834). Nonetheless, in all sessions, the main effect of category was significant (Session 1: *F* (2, 56) = 17.15, *p* < .001, ƞ^2^_G_ = .0120; Session 2: *F* (2, 54) = 12.33, *p* < .001, ƞ^2^_G_ = .0130; Session 3: *F* (1.55, 40.28) = 15.95, *p* < .001, ƞ^2^_G_ = .0150). A significant block *×* category interaction was only observed in the first session [*F* (2, 56) = 3.81, *p* = .028, ƞ^2^_G_ = .0139] but not in the second [*F* (1.32, 35.52) = .74, *p* = .430] or the third sessions [*F* (2, 52) = .94, *p* = .398]. Post hoc pairwise comparisons were only conducted for the first session since the block *×* category interaction was significant only in the first session. This revealed a significant increase in accuracy for lookalikes (*t*(28) = 2.36, *p* = .025, 95% CI [.006, .085]), whereas there was no significant change for animals (*t*(28) = *−*1.19, *p* = .244, 95% CI [*−*.064, .017]) and objects (*t*(28) = 1.54, *p* = .135, 95% CI [*−*.007, .046]).

### S1.2 Longitudinal Analysis

#### Reaction Time (Figure S1d)

One participant was excluded from the analysis based on the Z-score criterion. We conducted a 3 (session) *×* 3 (category) repeated-measures ANOVA on RT from extra before blocks across three sessions to examine the longitudinal effects of training. A significant main effect of session [*F* (2, 56) = 8.99, *p* < .001, ƞ^2^_G_ = .0134] and category [*F* (2, 56) = 26.41, *p* < .001, ƞ^2^_G_ = .0180] was found, although the session *×* category interaction did not reach statistical significance [*F* (4, 112) = 1.61, *p* = .177]. As the session *×* category interaction was not significant, no pairwise comparisons were performed.

#### Slowness (Figure S2d)

Two participants were excluded from the analysis based on the Z-score criterion. A 3 (session) *×* 3 (category) repeated-measures ANOVA demonstrated a significant main effect of session [*F* (1.59, 42.95) = 64.14, *p* < .001, ƞ^2^_G_ = .026] and category [*F* (2, 54) = 20.05, *p* < .001, ƞ^2^_G_ = .0180], as well as a significant session *×* category interaction [*F* (2.78, 74.93) = 6.40, *p* < .001, ƞ^2^_G_ = .026]. Thus, the reduction in the number of slow answers across sessions differed by category. Post hoc pairwise comparisons revealed that slowness decreased significantly from Session 1 to Session 2 (animals: *t*(27) = 7.61, *p* < .001, 95% CI [.041, .071]; lookalikes: *t*(27) = 7.68, *p* < .001, 95% CI [.101, .174]; objects: *t*(27) = 5.07, *p* < .001, 95% CI [.055, .129]). There was also a significant decrease in slowness from Session 1 to Session 3 (animals: *t*(27) = 6.65, *p* < .001, 95% CI [.037, .071]; lookalikes: *t*(27) = 7.51, *p* < .001, 95% CI [.100, .175]; objects: *t*(27) = 4.12, *p* = .001, 95% CI [.044, .131]) for all categories. However, none of the categories showed a significant difference between Session 2 and 3.

**Figure S3.**
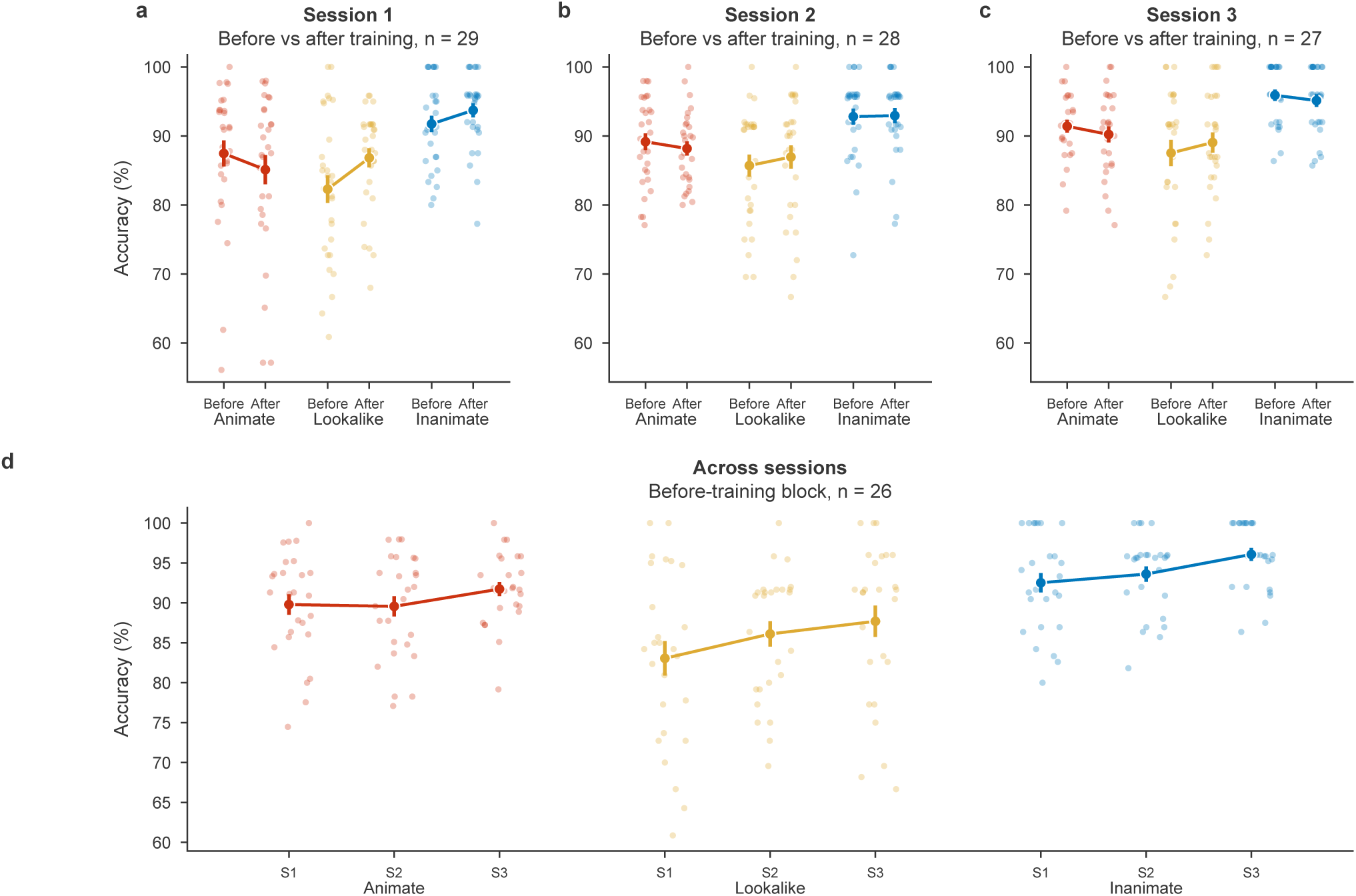
Accuracy during behavioural training: the proportion of correct responses among answered trials (too-slow trials are not scored). (a–c) Extra before and extra after blocks (fixed display time of 600 ms) of Sessions 1–3, for animals (animate), lookalikes and objects (inanimate). (d) Extra before block of each session. Dots show individual participants; filled circles and error bars show the group mean and standard error of the mean. Participants excluded by the Z-score criterion are not shown, and each panel title gives the number of participants included.s

#### Accuracy (Figure S3d)

Four participants were excluded from the analysis based on the Z-score criterion. A 3 (session) *×* 3 (category) repeated-measures ANOVA revealed that there was a significant main effect of session [*F* (2, 50) = 5.71, *p* = .006, ƞ^2^_G_ = .055] and category [*F* (2, 50) = 35.47, *p* < .001, ƞ^2^_G_ = .0360]. However, no significant session *×* category interaction was observed [*F* (2.78, 69.51) = .50, *p* = .667]. As the main effect of session was significant, post hoc pairwise comparisons between sessions (collapsed across categories; Bonferroni-corrected *p*-values) demonstrated that there was a significant increase in accuracy from Session 1 to Session 3 (*t*(25) = *−*3.25, *p* = .010, 95% CI [*−*.055, *−*.012]). However, no significant increase was observed for accuracy between Session 1 and Session 2 (*t*(25) = *−*1.22, *p* = .701, 95% CI [*−*.035, .009]) and Session 2 and Session 3 (*t*(25) = *−*2.28, *p* = .095, 95% CI [*−*.039, *−*.002]).

## References

Baeck, A., Maes, K., Van Meel, C., & Op de Beeck, H. (2016). The transfer of object learning after training with multiple exemplars. Frontiers in Psychology, 7. 10.3389/fpsyg.2016.01386

Baeck, A., Windey, I., & Op de Beeck, H. (2012). The transfer of object learning across exemplars and their orientation is related to perceptual similarity. Vision Research, 68, 40–47. 10.1016/j.visres.2012.06.023

Bang, J. W., Hamilton-Fletcher, G., & Chan, K. C. (2023). Visual plasticity in adulthood: Perspectives from hebbian and homeostatic plasticity. The Neuroscientist, 29 (1), 117–138. 10.1177/10738584211037619

Bracci, S., & Op de Beeck, H. (2016). Dissociations and associations between shape and category representations in the two visual pathways. Journal of Neuroscience, 36 (2), 432–444. 10.1523/JNEUROSCI.2314-15.2016

Bracci, S., & Op de Beeck, H. (2023). Understanding human object vision: A picture is worth a thousand representations. Annual Review of Psychology, 74, 113–135. 10.1146/annurev-psych-032720-041031

Bracci, S., Ritchie, J. B., Kalfas, I., & Op de Beeck, H. (2019). The ventral visual pathway represents animal appearance over animacy, unlike human behavior and deep neural networks. The Journal of Neuroscience, 39 (33), 6513–6525. 10.1523/JNEUROSCI.1714-18.2019

Bracci, S., Ritchie, J. B., & Op de Beeck, H. (2017). On the partnership between neural representations of object categories and visual features in the ventral visual pathway. Neuropsychologia, 105, 153–164. 10.1016/j.neuropsychologia.2017.06.010

Byers, A., & Serences, J. T. (2014). Enhanced attentional gain as a mechanism for generalized perceptual learning in human visual cortex. Journal of Neurophysiology, 112 (5), 1217–1227. 10.1152/jn.00353.2014

Carlson, T., Tovar, D. A., Alink, A., & Kriegeskorte, N. (2013). Representational dynamics of object vision: The first 1000 ms. Journal of Vision, 13 (10), 1–19. 10.1167/13.10.1

Carlson, T. A., Ritchie, J. B., Kriegeskorte, N., Durvasula, S., & Ma, J. (2014). Reaction time for object categorization is predicted by representational distance. Journal of Cognitive Neuroscience, 26 (1), 132–142. 10.1162/jocna00476

Chen, C.-Y., Leys, G., Bracci, S., & Op de Beeck, H. (2023). The representational dynamics of the animal appearance bias in human visual cortex are indicative of fast feedforward processing. Imaging Neuroscience, 1, imag-1–00006. 10.1162/imaga00006

Cheng, Y.-A., Sanayei, M., Chen, X., Jia, K., Li, S., Fang, F., Watanabe, T., Thiele, A., & Zhang, R.-Y. (2025). A neural geometry approach comprehensively explains apparently conflicting models of visual perceptual learning. Nature Human Behaviour, 9 (5), 1023–1040. 10.1038/s41562-025-02149-x

Cichy, R. M., Khosla, A., Pantazis, D., Torralba, A., & Oliva, A. (2016). Comparison of deep neural networks to spatio-temporal cortical dynamics of human visual object recognition reveals hierarchical correspondence. Scientific Reports, 6, 27755. 10.1038/srep27755

Cichy, R. M., Pantazis, D., & Oliva, A. (2014). Resolving human object recognition in space and time. Nature Neuroscience, 17 (3), 455–462. 10.1038/nn.3635

Clarke, A., Pell, P. J., Ranganath, C., & Tyler, L. K. (2016). Learning warps object representations in the ventral temporal cortex. Journal of Cognitive Neuroscience, 28 (7), 1010–1023. 10.1162/jocna00951

Cohen, L., Dehaene, S., Naccache, L., Lehéricy, S., Dehaene-Lambertz, G., Hénaff, M.-A., & Michel, F. (2000). The visual word form area: Spatial and temporal characterization of an initial stage of reading in normal subjects and posterior split-brain patients. Brain, 123 (2), 291–307. 10.1093/brain/123.2.291

Cohen, L., Lehéricy, S., Chochon, F., Lemer, C., Rivaud, S., & Dehaene, S. (2002). Language-specific tuning of visual cortex? functional properties of the visual word form area. Brain, 125 (5), 1054–1069. 10.1093/brain/awf094

Contini, E. W., Wardle, S. G., & Carlson, T. A. (2017). Decoding the time-course of object recognition in the human brain: From visual features to categorical decisions. Neuropsychologia, 105, 165–176. 10.1016/j.neuropsychologia.2017.02.013

Delorme, A., & Makeig, S. (2004). Eeglab: An open source toolbox for analysis of single-trial eeg dynamics including independent component analysis. Journal of Neuroscience Methods, 134 (1), 9–21. 10.1016/j.jneumeth.2003.10.009

DiCarlo, J. J., Zoccolan, D., & Rust, N. C. (2012). How does the brain solve visual object recognition? Neuron, 73 (3), 415–434. 10.1016/j.neuron.2012.01.010

Downing, P. E., Jiang, Y., Shuman, M., & Kanwisher, N. (2001). A cortical area selective for visual processing of the human body. Science, 293 (5539), 2470–2473. 10.1126/science.1063414

Duyck, S., Costantino, A. I., Bracci, S., & Op de Beeck, H. (2024). A computational deep learning investigation of animacy perception in the human brain. Communications Biology, 7 (1), 1718. 10.1038/s42003-024-07415-8

Epstein, R., & Kanwisher, N. (1998). A cortical representation of the local visual environment. Nature, 392 (6676), 598–601. 10.1038/33402

Ester, E. F., Sprague, T. C., & Serences, J. T. (2020). Categorical biases in human occipitoparietal cortex. The Journal of Neuroscience, 40 (4), 917–931. 10.1523/JNEUROSCI.2700-19.2019

Folstein, J., Palmeri, T. J., Van Gulick, A. E., & Gauthier, I. (2015). Category learning stretches neural representations in visual cortex. Current Directions in Psychological Science, 24 (1), 17–23. 10.1177/0963721414550707

Gergely, G., & Csibra, G. (2003). Teleological reasoning in infancy: The näıve theory of rational action. Trends in Cognitive Sciences, 7 (7), 287–292. 10.1016/S1364-6613(03)00128-1

Gergely, G., Ńadasdy, Z., Csibra, G., & Báró, S. (1995). Taking the intentional stance at 12 months of age. Cognition, 56 (2), 165–193. 10.1016/0010-0277(95)00661-H

Golan, T., Raju, P. C., & Kriegeskorte, N. (2020). Controversial stimuli: Pitting neural networks against each other as models of human cognition. Proceedings of the National Academy of Sciences, 117 (47), 29330–29337. 10.1073/pnas.1912334117

Gorgolewski, K. J., Auer, T., Calhoun, V. D., Craddock, R. C., Das, S., Duff, E. P., Flandin, G., Ghosh, S. S., Glatard, T., Halchenko, Y. O., Handwerker, D. A., Hanke, M., Keator, D., Li, X., Michael, Z., Maumet, C., Nichols, B. N., Nichols, T. E., Pellman, J., . . . Poldrack, R. A. (2016). The brain imaging data structure, a format for organizing and describing outputs of neuroimaging experiments. Scientific Data, 3, 160044. 10.1038/sdata.2016.44

Grill-Spector, K., & Weiner, K. S. (2014). The functional architecture of the ventral temporal cortex and its role in categorization. Nature Reviews Neuroscience, 15 (8), 536–548. 10.1038/nrn3747

Grootswagers, T., Robinson, A. K., Shatek, S. M., & Carlson, T. A. (2019). Untangling featural and conceptual object representations. NeuroImage, 202, 116083. 10.1016/j.neuroimage.2019.116083

Grootswagers, T., Wardle, S. G., & Carlson, T. A. (2017). Decoding dynamic brain patterns from evoked responses: A tutorial on multivariate pattern analysis applied to time series neuroimaging data. Journal of Cognitive Neuroscience, 29 (4), 677–697. 10.1162/jocna01068

Heider, F., & Simmel, M. (1944). An experimental study of apparent behavior. The American Journal of Psychology, 57 (2), 243–259. 10.2307/1416950

Hofrichter, R., Siddiqui, H., Morrisey, M. N., & Rutherford, M. D. (2021). Early attention to animacy: Change-detection in 11-month-olds. Evolutionary Psychology, 19 (2), 14747049211028220. 10.1177/14747049211028220

Jozwik, K. M., Najarro, E., van den Bosch, J. J. F., Charest, I., Cichy, R. M., & Kriegeskorte, N. (2022). Disentangling five dimensions of animacy in human brain and behaviour. Communications Biology, 5 (1), 1247. 10.1038/s42003-022-04194-y

Kaiser, D., Azzalini, D. C., & Peelen, M. V. (2016). Shape-independent object category responses revealed by meg and fmri decoding. Journal of Neurophysiology, 115 (4), 2246–2250. 10.1152/jn.01074.2015

Kanwisher, N., McDermott, J., & Chun, M. M. (1997). The fusiform face area: A module in human extrastriate cortex specialized for face perception. Journal of Neuroscience, 17 (11), 4302–4311. 10.1523/JNEUROSCI.17-11-04302.1997

Kato, M., & Mugitani, R. (2015). Pareidolia in infants. PLoS ONE, 10 (2), e0118539. 10.1371/journal.pone.0118539

Keefe, B. D., Gouws, A. D., Sheldon, A. A., Vernon, R. J. W., Lawrence, S. J. D., McKeefry, D. J., Wade, A. R., & Morland, A. B. (2018). Emergence of symmetry selectivity in the visual areas of the human brain: Fmri responses to symmetry presented in both frontoparallel and slanted planes. Human Brain Mapping, 39 (10), 3813–3826. 10.1002/hbm.24211

Khaligh-Razavi, S.-M., & Kriegeskorte, N. (2014). Deep supervised, but not unsupervised, models may explain it cortical representation. PLoS Computational Biology, 10 (11), e1003915. 10.1371/journal.pcbi.1003915

Konkle, T., & Caramazza, A. (2013). Tripartite organization of the ventral stream by animacy and object size. Journal of Neuroscience, 33 (25), 10235–10242. 10.1523/jneurosci.0983-13.2013

Kriegeskorte, N., Mur, M., Ruff, D. A., Kiani, R., Bodurka, J., Esteky, H., Tanaka, K., & Bandettini, P. A. (2008). Matching categorical object representations in inferior temporal cortex of man and monkey. Neuron, 60 (6), 1126–1141. 10.1016/J.NEURON.2008.10.043

Long, B., Störmer, V. S., & Alvarez, G. A. (2017). Mid-level perceptual features contain early cues to animacy. Journal of Vision, 17 (6), 20. 10.1167/17.6.20

Long, B., Yu, C.-P., & Konkle, T. (2018). Mid-level visual features underlie the high-level categorical organization of the ventral stream. Proceedings of the National Academy of Sciences, 115 (38), E9015–E9024. 10.1073/pnas.1719616115

Maniquet, T., Op de Beeck, H., & Costantino, A. I. (2025). Recurrent issues with deep neural network models of visual recognition. Scientific Reports, 15 (1). 10.1038/s41598-025-20245-w

Mayhew, S. D., Li, S., Storrar, J. K., Tsvetanov, K. A., & Kourtzi, Z. (2010). Learning shapes the representation of visual categories in the aging human brain. Journal of Cognitive Neuroscience, 22 (12), 2899–2912. 10.1162/jocn.2010.21415

Nairne, J. S., VanArsdall, J. E., Pandeirada, J. N. S., Cogdill, M., & LeBreton, J. M. (2013). Adaptive memory: The mnemonic value of animacy. Psychological Science, 24 (10), 2099–2105. 10.1177/0956797613480803

Naselaris, T., Stansbury, D. E., & Gallant, J. L. (2012). Cortical representation of animate and inanimate objects in complex natural scenes. Journal of Physiology-Paris, 106 (5–6), 239–249. 10.1016/j.jphysparis.2012.02.001

O’Bryan, S. R., Jung, S., Mohan, A. J., & Scolari, M. (2024). Category learning selectively enhances representations of boundary-adjacent exemplars in early visual cortex. Journal of Neuroscience, 44 (3), e1039232023. 10.1523/JNEUROSCI.1039-23.2023

Oostenveld, R., Fries, P., Maris, E., & Schoffelen, J.-M. (2011). Fieldtrip: Open source software for advanced analysis of meg, eeg, and invasive electrophysiological data. Computational Intelligence and Neuroscience, 2011, 156869. 10.1155/2011/156869

Oosterhof, N. N., Connolly, A. C., & Haxby, J. V. (2016). Cosmomvpa: Multi-modal multivariate pattern analysis of neuroimaging data in matlab/gnu octave. Frontiers in Neuroinformatics, 10, 27. 10.3389/fninf.2016.00027

Peirce, J., Gray, J. R., Simpson, S., MacAskill, M., Höchenberger, R., Sogo, H., Kastman, E., & Lindeløv, J. K. (2019). Psychopy2: Experiments in behavior made easy. Behavior Research Methods, 51 (1), 195–203. 10.3758/s13428-018-01193-y

Pernet, C. R., Appelhoff, S., Gorgolewski, K. J., Flandin, G., Phillips, C., Delorme, A., & Oostenveld, R. (2019). Eeg-bids, an extension to the brain imaging data structure for electroencephalography. Scientific Data, 6 (1), 103. 10.1038/s41597-019-0104-8

Pillet, I., Cerrahoglu, B., Philips, R. V., Dumoulin, S., & Op de Beeck, H. (2024). A 7T fMRI investigation of hand and tool areas in the lateral and ventral occipitotemporal cortex. PLOS ONE, 19 (11), e0308565. 10.1371/journal.pone.0308565

Premack, D., & Woodruff, G. (1978). Does the chimpanzee have a theory of mind? Behavioral and Brain Sciences, 1 (4), 515–526. 10.1017/S0140525X00076512

Proklova, D., Kaiser, D., & Peelen, M. V. (2019). MEG sensor patterns reflect perceptual but not categorical similarity of animate and inanimate objects. NeuroImage, 193, 167–177. 10.1016/j.neuroimage.2019.03.028

R Core Team. (2025). R: A language and environment for statistical computing. R Foundation for Statistical Computing. Vienna, Austria. https://www.R-project.org/

Rakison, D. H., & Poulin-Dubois, D. (2001). Developmental origin of the animate–inanimate distinction. Psychological Bulletin, 127 (2), 209–228. 10.1037/0033-2909.127.2.209

Ritchie, J. B., Tovar, D. A., & Carlson, T. A. (2015). Emerging object representations in the visual system predict reaction times for categorization. PLoS Computational Biology, 11 (6), e1004316. 10.1371/journal.pcbi.1004316

Ritchie, J. B., Zeman, A. A., Bosmans, J., Sun, S., Verhaegen, K., & Op de Beeck, H. (2021). Untangling the animacy organization of occipitotemporal cortex. Journal of Neuroscience, 41 (33), 7103–7119. 10.1523/JNEUROSCI.2628-20.2021

Rostami, F., Spriet, C., Op de Beeck, H., Hochmann, J.-R., & Papeo, L. (2026). Hard-to-beat animacy perception: EEG evidence of accurate animate-inanimate distinction for ambiguous objects. bioRxiv. 10.64898/2026.06.15.732334

Serra, M. J., & DeYoung, C. M. (2023). The animacy advantage in memory occurs under self-paced study conditions, but participants’ metacognitive beliefs can deter it. Frontiers in Psychology, 14. 10.3389/fpsyg.2023.1164038

Serra, M. J., Pandeirada, J. N. S., & VanArsdall, J. E. (2024). Editorial: Animacy in cognition: Effects, mechanisms, and theories. Frontiers in Psychology, 15. 10.3389/fpsyg.2024.1508218

The MathWorks Inc. (2024). Matlab version: 24.2.0.2712019 (r2024b). The MathWorks Inc. Natick, Massachusetts, United States. https://www.mathworks.com

Townsend, J. T., & Ashby, F. G. (1978). Methods of modeling capacity in simple processing systems. In N. J. Castellan & F. Restle (Eds.), Cognitive theory (pp. 200–239, Vol. 3). Erlbaum.

Van Gulick, A. E., & Gauthier, I. (2014). The perceptual effects of learning object categories that predict perceptual goals. *Journal of Experimental Psychology: Learning*, Memory, and Cognition, 40 (5), 1307–1320. 10.1037/a0036822

Wardle, S. G., Taubert, J., Teichmann, L., & Baker, C. I. (2020). Rapid and dynamic processing of face pareidolia in the human brain. Nature Communications, 11 (1), 4518. 10.1038/s41467-020-18325-8

Watanabe, T., & Sasaki, Y. (2015). Perceptual learning: Toward a comprehensive theory. Annual Review of Psychology, 66, 197–221. 10.1146/annurev-psych-010814-015214

Yamins, D. L. K., Hong, H., Cadieu, C. F., Solomon, E. A., Seibert, D., & DiCarlo, J. J. (2014). Performance-optimized hierarchical models predict neural responses in higher visual cortex. Proceedings of the National Academy of Sciences, 111 (23), 8619–8624. 10.1073/pnas.1403112111

Yeh, L.-C., & Peelen, M. V. (2022). The time course of categorical and perceptual similarity effects in visual search. Journal of Experimental Psychology: Human Perception and Performance, 48 (10), 1069–1082. 10.1037/xhp0001034

